# Genomic signals of local adaptation in *Eleginops maclovinus* from Northern Chilean Patagonia

**DOI:** 10.1101/2024.12.20.629640

**Authors:** C. Eliza Claure, Garrett D. McKinney, J. Dellis Rocha, José M. Yáñez, Iván Pérez-Santos, Cristian B. Canales-Aguirre

## Abstract

Understanding the evolutionary mechanisms that shape the adaptive divergence across spatially heterogeneous environments is a challenging task for evolutionary ecologists. The Chilean marine Patagonia is a complex ecosystem with diverse geomorphology and physical-chemical oceanographic conditions. There is limited research evaluating the interactions between selective forces and environmental conditions in this area. This study focuses on identifying the genomic signals of local adaptation of the endemic marine fish, *Eleginops maclovinus* from Chilean North Patagonia. To achieve this goal, we used an environmental marine database (temperature, salinity, oxygen, phosphate nitrate and silicate concentration) with collected from 1995 to 2018 and 11,961 SNPs obtained from 246 individuals from 10 sampling locations across this area. We identified putative adaptive loci using ten bioinformatic software tools, where five were based on population genetic differentiation (PGD) and five based on the genotype-environment association (GEA). We identified 392 adaptative loci using PGD and 2,164 associated with at least one of the six environmental variables analyzed using GEA. A total of 131 loci were shared between the PGD and GEA approaches, of which 37 were associated with genes involved in the growth, metabolism and homeostasis. Then, we evaluated the variation of adaptive loci with environmental variables using polygenic scores and found significant correlations with temperature, salinity, and oxygen, indicating polygenic selection along environmental gradients. This study highlights how polygenic selection drives local adaptation in *Eleginops maclovinus* and underscores the value of integrating genomic and environmental data for conservation in the Patagonian ecosystem.

## 1 Introduction

The main aim of adaptive genomics is understanding the molecular basis of local adaptation in species experiencing spatially heterogeneous environments. Currently, detecting adaptative loci is possible using high-throughput sequencing technologies, which allows thousands of single-nucleotide polymorphisms (SNPs) distributed across the genome (Willette *et al*. 2014), to be genotyped in hundreds of individuals (Manel *et al*. 2016).

Restriction site-associated DNA sequencing (RAD-seq) method harness the high-throughput sequencing technology to genotype individuals (Miller *et al*. 2007; Andrews *et al*. 2016; McKinney *et al*. 2017a). Specifically, this method uses restriction enzymes that cut at specific motifs throughout the genome to reduce their complexity (Altshuler *et al*. 2000; Miller *et al*. 2007; Baird *et al*. 2008). The use of hundreds of thousands of RAD-seq markers for exhibiting the signatures of selection is increasingly applied to marine organisms, offering particular advantages for complex genomes in the absence of a reference sequence (Wang *et al*. 2015; Benestan *et al*. 2016; Cure *et al*. 2017; Dalongeville *et al*. 2018; Hoey & Pinsky 2018; Li *et al*. 2019; Segovia *et al*. 2020; O’Leary *et al*. 2021).

For identifying adaptative loci, there are two predominant genome scan approaches: *(i)* population genetic differentiation (PGD) and *(ii)* genotype-environment association (GEA). The approach based on population genetic differentiation (PGD) identifies loci putatively under selection by primarily comparing the genetic differentiation index (F_ST_) and other measures (e.g., inbreeding coefficient (F_IS_) and locus-specific homozygosity excess) for each locus against values predicted by a null model of neutral evolution. If natural selection favors one allele at a particular locus in some populations, F_ST_ at that locus will be large compared to loci in which among-population differences are the result of neutral demographic processes (François *et al*. 2016). Contrarily, approach based on genotype-environment association (GEA) search for associations between environmental variables and allele frequencies across a set of populations spread over a geographic space (Rellstab *et al*. 2015; Hoban *et al*. 2016). Alleles involved in local adaptation should occur at higher frequency where they increase fitness and be at lower frequency where they decrease fitness. Therefore, it may be possible to identify putative adaptative loci by scanning for alleles that show greater than average genetic differentiation among populations using GEA approach (Lewontin & Krakauer 1973; Beaumont 2005). These approaches are based on the premise that it is possible to distinguish between the genetic signals left by adaptative and neutral processes. However, algorithms, statistical models, and prior assumptions considerably differ and all of them are prone to false positive associations (de Villemereuil *et al*. 2014; Rellstab *et al*. 2015; de Villemereuil & Gaggiotti 2015; François *et al*. 2016). Combining PGD and GEA outcomes may provide an efficient strategy to detect signatures of local adaptation while reducing the number of false positives (de Villemereuil *et al*. 2014; Rellstab *et al*. 2015; Gagnaire *et al*. 2015; François *et al*. 2016).

North Patagonian ecosystems in southern Chile are constituted by extensive and interconnected system of fjords, channels, gulf, estuaries, and bays that are affected by physical regimes that may strongly modulate biological productivity (Pantoja *et al*. 2011; Iriarte *et al*. 2014). It extends approximately across 140,000 km^2^ from latitude ∼41.5°S (Reloncaví Fjord) to latitude ∼46.5°S (San Rafael Lagoon) (Rodrigo 2008). Oceanographically, Chilean Northern Patagonia is considered a transitional marine system, where the interaction between marine and fresh waters produces strong vertical and horizontal gradients in physical-chemical variables (Pickard 1971; Dávila *et al*. 2002; Iriarte *et al*. 2014; Pérez-Santos *et al*. 2014; Schneider *et al*. 2014). These characteristics make Patagonia an excellent natural laboratory to investigate how this seascape heterogeneity results in genomics features. Thus, integrating genomic and environmental data provides a powerful approach for uncovering the processes leading to adaptative divergence in ecologically heterogeneous and connected marine ecosystems. So far, studies focused on seascape genomics in Northern Patagonia are limited (Araneda *et al*. 2016; Canales-Aguirre *et al*. 2016, 2018; Delgado *et al*. 2019; Segovia *et al*. 2020; Canales-Aguirre *et al*. 2022).

*Eleginops maclovinus* (Cuvier & Valenciennes 1830), is a notothenioid fish (Perciformes) of the monotypic family Eleginopidae (Pequeño 1989). This marine fish is endemic to the coastal temperate and sub-Antarctic waters of South America with a distribution range along the Atlantic and Pacific Patagonian coasts. In the Atlantic Ocean, it is distributed from Uruguay (∼35°S) to the Beagle Channel (∼54°S), including the Falkland/Malvinas Islands (Gosztonyi & Ae 1974; Eastman 1993), while in the Pacific Ocean, from the Beagle Channel to Valparaíso (∼33°S) (Pequeño 1989). This species is a protandrous hermaphrodite with the highest fecundity (∼796 eggs g^−1^) (Brickle *et al*. 2005) among notothenioids (80-300 eggs g^−1^) (Kock 1992; North 2001). It is euryhaline and can tolerate a broad range of salinity concentrations (5 and 45 PSU), however, extreme salinities can produce dramatic metabolic responses to fuel osmoregulatory tissues (Vargas-Chacoff *et al*. 2014, 2015a). Furthermore, juveniles of *E. maclovinus* have been described having a large eurythermal capacity, with tolerance between 4 and 20°C (Vanella *et al*. 2012; Oyarzún *et al*. 2018; Lattuca *et al*. 2018). All these studies described above have been conducted under controlled conditions, they are focused on one or two environmental variables, and do not associate with genetic data. The latter leaves a gap on how loci under selection can shape the adaptative capacity of *E*. *maclovinus* populations considering the complex environmental variability in estuaries and fjords.

Studies about the geographic pattern of genetic diversity in *E*. *maclovinus* are limited to three (Ceballos *et al*. 2012, 2016; Canales-Aguirre *et al*. 2022). A phylogeographic research conducted by Ceballos et al. (2012) using mitochondrial DNA sequences for nine locations did not find any evidence for the structuring of genetic variation into an Atlantic and a Pacific group. Subsequently, using nine microsatellites it was revealed a low but significant level of genetic differentiation between Pacific and Atlantic populations and showed differences between locations in the Pacific coast (Ceballos *et al*. 2016). However, the authors acknowledge that a single mitochondrial locus or nine microsatellites may not be robust enough to allow them to recover significant subdivision among geographic regions. Recently, Canales-Aguirre *et al*. (2022) used 12,050 SNPs in five locations distributed across Northern Patagonia to identify neutral and adaptive population structure. However, the latter only included five locations, used a public environmental database Bio-ORACLE (Tyberghein *et al*. 2012; Assis *et al*. 2018) which is not accurate for fine-scale and fjords areas like Patagonia, and only three approaches to identify adaptive loci. In addition, its focus was to identify natural and adaptive loci for conservation and management purposes. To our knowledge, this is the first study to identify the signal of local adaptation using ten different bioinformatic approaches and six highly accurate environmental variables in populations of *E. maclovinus* along Northern Patagonia.

In this study, we applied RAD-seq sequencing to generate SNPs genotyped in 246 individuals of *E. maclovinus* collected in North Patagonia. Here, we expanded the sampling locations from Canales-Aguirre et al. (2022), increasing the number to ten locations to better capture the environmental heterogeneity of their North Patagonian seascape. Our aim was to detect SNPs putatively under selection using population genetic differentiation (PGD) and genotype-environment association (GEA) approaches. We tested associations between SNPs and six environmental variables (temperature, salinity concentration of oxygen, nitrate, phosphate and silicate) obtained from oceanographic data recorded between 1995 to 2018 in this area. We implemented a BLASTx search on putative adaptative SNPs identified by both PGD and GEA approaches to investigate genes with molecular functions potentially involved in local adaptation. Then, we evaluated the variation of adaptive loci using polygenic scores where we found significant correlations and supporting polygenic selection in North Patagonia.

## 2 Materials and Methods

### 2.1 Study area and environmental predictors

Chilean Patagonia is one of the most extended fjord regions in the world, located on the southeastern border of South America (Fig.1A), and composed by islands, peninsulas, fjords, and channels (Pantoja *et al*. 2011; Iriarte *et al*. 2014). This region has been affected by ice sheets calving into the ocean, retraction of the sea coast line and decrease in marine water temperatures since the Great Patagonian Glaciation (c. 1 Ma) during the Pleistocene (Rabassa *et al*. 2011). The Northern Patagonia is characterized by interactions between marine and fresh water, generating sharp vertical and horizontal salinity gradients (Dávila *et al*. 2002; Pérez-Santos *et al*. 2014; Schneider *et al*. 2014). The temperature shows a latitudinal gradient, which decreases southward until the Antarctic Polar Front (∼55°S) (Iriarte *et al*. 2014). The oxygen concentration in the fjords is the result of horizontal advection of adjacent well-oxygenated Subantarctic Waters (5–6 mL/L) representing the major source of oxygen in the deep layers of the inner seas of Patagonia (Silva & Vargas 2014). In recent years, dissolved oxygen (DO) in the fjords of Chilean Patagonia has decreased by 1–2%, potentially driven by nutrient accumulation and rising temperatures (Linford *et al*. 2023a; b, 2024). The concentrations of inorganic nutrients show a strong seasonal signal, with high nitrate and orthophosphate concentrations during winter, and lower values during spring, presumably caused by a severe increase in primary productivity when light availability in near-surface waters increases (Iriarte *et al*. 2007; González *et al*. 2011; Torres *et al*. 2014; Jacob *et al*. 2014). The concentration of silicic acid shows latitudinal and longitudinal gradient due to surface water mixing between nitrate-rich but Silicic acid-depleted oceanic subantarctic waters and Silicic acid-rich but nitrate depleted continental waters (Aracena *et al*. 2011; González *et al*. 2011; Torres *et al*. 2014).

**Figure 1.**
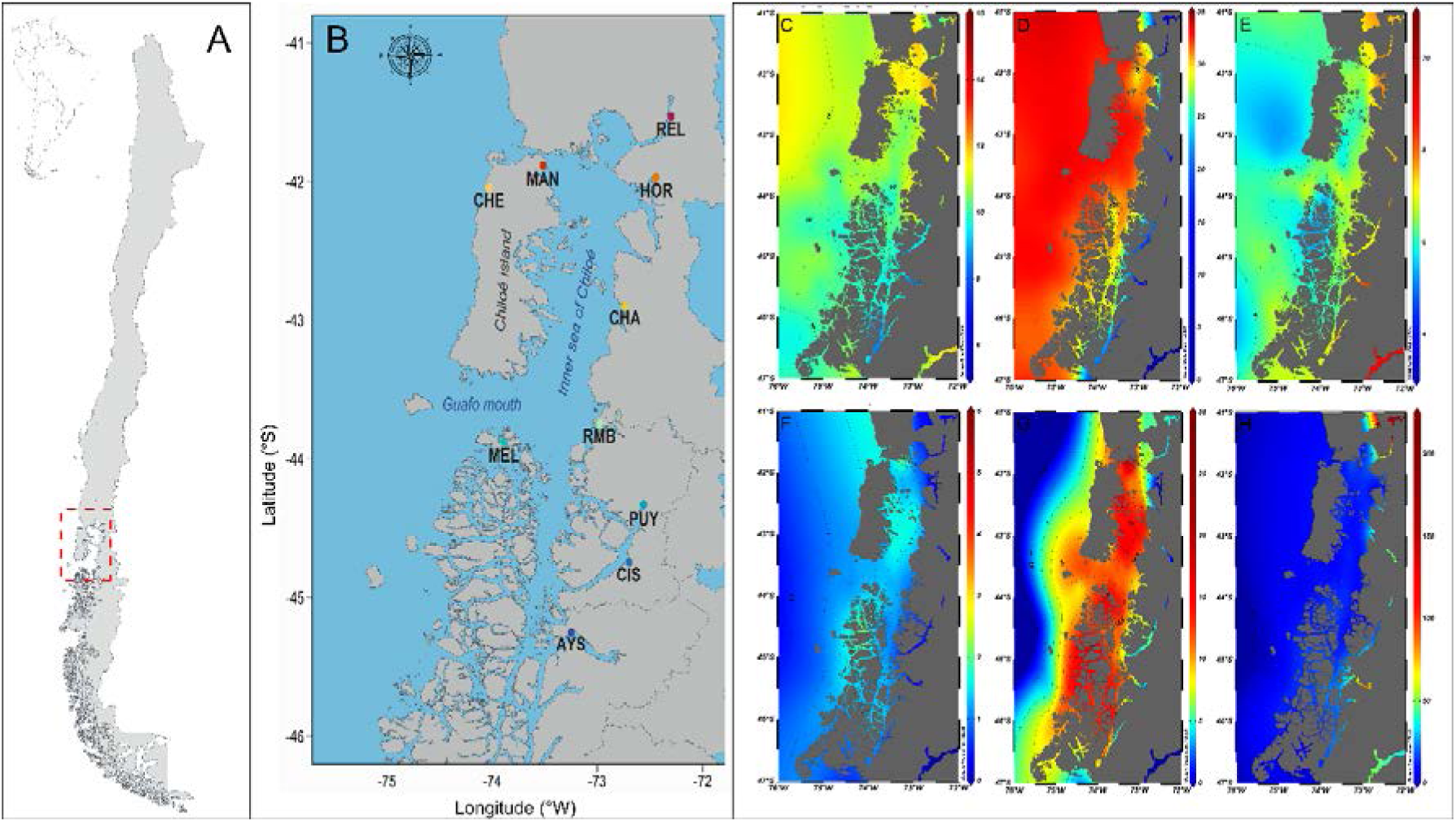
Geographic area and environmental variables in used. Map of sampling locations in South America. B) Sampling site locations on North Patagonian fjord ecosystems in southern Chile REL: Reloncaví Estuary, MAN: Manao, HOR: Hornopirén, CHE: Chepu, CHA: Chaitén, RMB: Raúl Marín Balmaceda, MEL: Melinka Island, PUY: Puyuhuapi, CIS: Port Cisnes, AYS: Port Aysén: C) Mean surface temperature (°C) D) Mean surface salinity (g/Kg); E) Mean surface concentration oxygen (µM); F) Mean surface nitrate (µM); G) Mean surface phosphate(µM); H) Mean surface silicate (µM).

We compiled an environmental database using data collected through the CIMAR Program of the National Oceanographic Committee of Chile (CONA), from CIMAR 1 (1995) to CIMAR 24 (2018). The database comprises 1,195 sampling stations located along the inner sea of Chiloé Island, estuaries, channels, and fjords from the Chilean Patagonia sampled in spring and winter (http://www.shoa.cl/n_cendhoc/productos/reporte_datos.php) (Fig. S1). At each sampling station, environmental variables were measured from the surface to the maximum possible depth. Considering that traditional databases lack high-resolution data near the coasts (e.g., Bio-Oracle, World Ocean Atlas, Aqua/MODIS), we highlight the use of information obtained by CIMAR as an effective alternative to detect the environmental heterogeneity of Northern Patagonia. Our database compiled contains six environmental predictors: conservative temperature^1^ (°C), absolute salinity^2^ (g Kg^−1^), oxygen (mL L^−1^), nitrate (µM), phosphate (µM) and silicate (µM) concentrations. We used Ocean Data View v.5.4.0. (Schlitzer 2002) for data visualization. We estimated the mean, minimum, maximum and range of these environmental variables in each of the ten locations for a water column until 100 m depth using the entire period of the database (1995 – 2018) (Table S1). These latter, considering that *E. maclovinus* inhabits in coastal and estuarine areas at depths >40 m, and the spawning occurs in spring season (Brickle *et al*. 2005)

### 2.2 Sampling and genotyping

Between November 2018 and April 2019, we sampled 246 individuals of *E. maclovinus* from ten locations throughout the North Patagonian fjord ecosystems in southern Chile (Fig.1B). Five of the ten sampling locations included in this study were previously analyzed by Canales-Aguirre *et al*. (2022). Sampling, DNA extraction, RAD-sequencing library preparation, sequencing with Illumina technology and pipeline of bioinformatic analyses were performed following procedures described by Canales-Aguirre *et al*. (2022).

Loci and samples were filtered iteratively by initially removing poor-quality data, recalculating missing data proportions, and applying increasingly stringent thresholds, following McKinney et al. (2020). The first part of the filtration process consisted in excluding SNPs or individuals based on the following criteria *(i)* minor allele frequencies (MAF) per sampling location, *(ii)* genotype rate per locus, *(iii)* genotype rate per sample. Initially, filtration criterion was MAF ≤ 0.05, genotype rate threshold of 50% per locus and sample. Subsequently, MAF≤ 0.1, genotype rate threshold of 90% per locus and sample. Due to gene duplication events described to have affected the evolution of the notothenioid genome (Xu *et al*. 2008; Chen *et al*. 2008, 2019), *HDplot* (McKinney *et al*. 2017b) was used to identify paralogs, which were removed from further analysis (parameter flags: H < 0.6; |D| < 5). We removed SNPs in Hardy-Weinberg disequilibrium (p-value < 0.05) if they deviated in three or more locations to avoid potential issues such as inbreeding, population stratification, and genotyping errors in the identified SNPs (Wigginton *et al*. 2005). For reads with more than one SNP, we only kept those with the highest F_ST_, to reduce the influence from linked loci in the results. The resulting filtered (.vcf) file was converted into the corresponding file formats for the subsequent analyses by using *PGDSpider* v. 2.1.1.5. (Lischer & Excoffier 2012).

### 2.3 Detecting putative loci under selection

We employed ten different software programs to identify putative adaptive loci: five based on population genetic differentiation (PGD) and five on genotype-environment association (GEA). We combined multiple approaches to minimize false positive rates and maximize our chances of detecting genomic signals of local adaptation (Rellstab *et al*. 2015; François *et al*. 2016). For the PGD approach, we used *fsthet* v.1.0.1 (Flanagan & Jones 2017), *BayeScan* v.2.1 (Foll & Gaggiotti 2008), *OutFLANK* v.0.2 (Whitlock & Lotterhos 2015), *PCAdapt* v.4.3.3 (Luu *et al*. 2017; Privé *et al*. 2020), and *Arlequin* v.3.5.2.2 (Excoffier & Lischer 2010) (Table 1). For the GEA approach, we used six environmental variables and five software programs: *Latent Factors Mixed Model (LFMM)* (Frichot *et al*. 2013), *BayeScEnv* (de Villemereuil & Gaggiotti 2015), *Redundancy Analysis (RDA).*(Hair *et al*. 1995; Zuur *et al*. 2010; Capblancq *et al*. 2018; Capblancq & Forester 2021)*, Moran Spectral Outlier Detection (MSOD)* (Wagner *et al*. 2017), and *Sam*β*ada* (Stucki *et al*. 2017; Duruz *et al*. 2019) (Table 2). In all programs, a 5% False Discovery Rate and respective p-values or q-values were applied to detect putative adaptive loci. We employed *UpSet* diagrams (Lex *et al*. 2014) to visually compare the results from the ten software programs, identifying loci flagged as outliers by at least two programs as putative adaptive loci. For subsequent analyses, loci were classified into two categories: putative adaptive loci (under directional selection) and putative neutral loci, excluding loci under balancing selection.

**Table 1.** Description and parameters used for software with the PGD approach.

| Software | Description | Parameters | References |
| --- | --- | --- | --- |
| i) *fsthet* | *fsthet* identifies loci with extreme $F_{ST}$ values by calculating smoothed quantiles for the $F_{ST}$-heterozygosity distribution, using its own empirical distribution instead of a null model due to changes in population structure. | We used a confidence level of 95% to detect $F_{ST}$ outliers | (Flanagan & Jones 2017) |
| ii) *BayeScan* | *BayeScan* estimates the probability of selection for each locus using a Bayesian test that distinguishes between locus-specific selection effects ($\alpha$) and population-specific demography effects ($\beta$), by measuring the discordance between global and population-specific allele frequencies. | We ran it with default parameters: 20 pilot runs of 5,000 iterations followed by 50,000 iterations with an additional burn-in of 50,000 and prior odd of 10. | (Foll & Gaggiotti 2008) |
| iii) *OutFLANK* | OutFLANK calculates a neutral distribution of $F_{ST}$ values and assigns q-values to each locus to detect adaptive loci influenced by selection, without assuming a specific population's demographic history, reducing false positives. | We ran it with default parameters: LeftTrimFraction = 0.05, RightTrimFraction = 0.05, Hmin = 0.1. | (Whitlock & Lotterhos 2015) |
| iv) *PCAdapt* | *PCAdapt* performs a Principal Component Analysis (PCA) and computes the p-values for each locus to detect putative adaptative loci. | The most appropriate number of clusters K was chosen by testing K=1:10. We detected outliers using K=2 and p-values < 0.05 associated with Mahalanobis distances calculated with the q-value package. | (Dabney *et al.* 2010; Luu *et al.* 2017) |
| v) *Arlequin* | *Arlequin* calculates observed heterozygosity ($H_O$) to create a null distribution of $F_{ST}$ values and associated p-values for each locus. | This was executed under a hierarchical island model with 20,000 simulations and 100 demes, which implements FDIST2 methodology. | (Excoffier & Lischer 2010) |

**Table 2.** Description and parameters used for software with the GEA approach.

| Software | Description | Parameters | References |
| --- | --- | --- | --- |
| i) Latent factors mixed model (*LFMM*) | *LFMM* is a hierarchical Bayesian mixed model, which uses latent factors to correct population structure | We estimated the number of ancestral populations K before the analysis testing from K 1 to 10 using the LEA v.3.2.0 R package and determined K from the cross-entropy criteria and Cattell's | (Cattell 1966; Devlin & Roeder 1999; Frichot *et al.* |
|  | while fitting a linear regression between allele frequencies and environmental variables. | rule from the sNMF output. LFMM analyses were conducted separately for each environmental variable, with 10,000 iterations, and burning of 5,000 and five replicates. | 2013, 2014; Frichot & François 2015) |
| ii) *BayeScEnv* | *BayeScEnv* detects putative adaptive loci by identifying loci that show large positive $F_{ST}$ values (outside the neutral model $F_{ST}$ distribution) that are significantly correlated with environmental variables. | We used 20 pilot runs, with 50,000 iterations each and an additional burn-in of 50,000 iterations for each of the six standardized environmental variables. | (de Villemereuil & Gaggiotti 2015). |
| iii) *Redundancy Analysis (RDA)* | *Redundancy Analysis (RDA)* is a multivariate ordination method to detect loci putatively under selection based on correlations with environmental variables. | We calculated the variance inflation factor (vif.cca function) to exclude variables with a value $\geq 5$, and conducted linear regressions between allele frequencies and environmental variables at each locus. Using PCA, we produced ordination axes, identified putative adaptive loci ($\pm 3$ SD from the mean), and evaluated the correlation using the Pearson correlation coefficient. | (Hair *et al.* 1995; Zuur *et al.* 2010; Rellstab *et al.* 2015; Forester *et al.* 2018) |
| iv) *Moran spectral outlier detection (MSOD)* | *MSOD* uses Moran eigenvector maps (MEM) to quantify the distribution of allele frequencies across a range of spatial scales represented by MEM spatial eigenvectors | We compared the power spectrum of each locus to the average power spectrum to identify outliers using z-scores and a cut-off of 0.05. Subsequently, we applied Moran spectral randomization (MSR) with 999 permutations using adespatial. | (Dray *et al.* 2006, 2021; Wagner & Dray 2015; Wagner *et al.* 2017) |
| v) *Samβada* | *Samβada* is a spatial approach that uses multiples univariate logistic regression models to identify locus-environment associations and at the same time measures spatial autocorrelation | We coded individuals for the presence or absence of each SNP allele and measured the association with environmental parameters across sites using multiple logistic regression. P-values from these regressions were calculated via a Wald Score Test and compared to a $\chi^2$ distribution with one degree of freedom. | (Wald 1943; Stucki *et al.* 2017; Duruz *et al.* 2019) |

### 2.4 Genetic diversity and population structure

For the adaptative and neutral datasets, we determined, described and compared overall and population-specific genetic diversity. The expected heterozygosity (H_E_), observed heterozygosity (H_O_) and inbreeding coefficient (F_IS_) were calculated for each locus and locations using *Hierfstat* v 0.04.10 R package (Goudet 2005). We assessed the genetic differentiation with pairwise F_ST_ (Weir & Cockerham 1984) comparisons between locations using the R package *StAMPP* v.1.6.2. (Pembleton *et al*. 2013) and the statistical significances were determined using 10,000 permutations.

We used two approaches to investigate the genetic structure: Bayesian clustering program *Structure* v.2.3.4. (Pritchard *et al*. 2000) and discriminant analysis of principal components (DAPC) (Jombart *et al*. 2010) using r package *adegenet* v.2.1.5. (Jombart 2008; Jombart & Ahmed 2011). *Structure* is a Bayesian method, which uses a Markov chain Monte Carlo (MCMC) algorithm based on the Gibbs sampler algorithm to identify the number of putative genetic clusters. This method assumes Hardy-Weinberg equilibrium (HWE) and linkage equilibrium among loci in sample population individuals (Pritchard *et al*. 2000). We conducted the analysis using ten replicates per K value for K=1 to K=10, with 10,000 burn-in steps and 100,000 MCMC-sampling steps. The runs were collected, sorted and merged and functions in the R package *pophelper* (Francis 2017) to produce assignment barplots. The most likely number of genetic clusters (K) was chosen using the ΔK method (Evanno *et al*. 2005). The DAPC method identifies and describes clusters of genetically related individuals from genomic datasets and allows the optimal visualization of between-population differentiation in multivariate space (Jombart *et al*. 2010). First, we carried out a principal component analysis (PCA), and used the principal components (PCs) thus produced as synthetic variables for a discriminant analysis, as outlined in Jombart et al. (2010). The optimal number of principle components was determined by cross validation (*xvalDapc* function (Jombart *et al*. 2010)) to avoid overfitting.

We also conducted isolation by distance (IBD) analyses to test whether genetic distance increased with geographic distance with adaptative and neutral data. We assessed the correlation between the linearized pairwise F_ST_ matrix and the Euclidean (geographic) distance matrix using a Mantel-test as implemented in the R-package *vegan* (Oksanen *et al*. 2016). Statistical significance was assessed with 999 permutations of the data. To test whether the correlation between genetic and geographic distances is a result of a continuous or distant patchy cline of genetic differentiation, local densities of distances to disentangle both processes were plotted using 2-dimensional kernel density estimation (*kde2d* function) of *MASS* R-package (Venables & Ripley 2013).

### 2.5 Additive polygenic scores (APS)

We assessed cumulative adaptive genetic variation by calculating additive polygenic scores (APS) associated with each environmental variable (Gagnaire & Gaggiotti 2016; Babin *et al*. 2017). For this analysis, we prioritized three biologically important environmental variables: annual maximum temperature (°C), annual maximum salinity (g/Kg), and annual minimum oxygen concentration (µM) (Holliday 1969; Nikinmaa & Rees 2005; Brian *et al*. 2008; Vanella *et al*. 2012; Vargas-Chacoff *et al*. 2014). We prioritized the maximum values of temperature and salinity considering the effect that climate change has on them. According to the Community Climate System Model 3.0 (CCSM3) of the National Center for Atmospheric Research (NCAR), a 2.5°C increase in the temperature of Chilean coastal areas is predicted by 2055 (Silva *et al*. 2016). Similarly, the Center for Oceanographic Research in the Eastern South Pacific (COPAS) detected an increase of 0.25 in salinity in the upper layers of the central-southern coast of Chile (Schneider *et al*. 2017)We also considered the minimum oxygen concentration, as hypoxic conditions negatively impact fish fitness (Richards 2009; Wu 2009). All these variables have an impact of the adaptive genetic diversity of several species (Pauls *et al*. 2013; Bernatchez 2016)..

We assessed the relationship between each of the 131 putative adaptative loci and each environmental variable. We used genotypes (0,1,2) to determine the score for each favored allele; if the slope is positive, and (2,1,0) if negative (Babin *et al*. 2017). We calculated APS by summing the scores of favored alleles associated with each environmental variable. Finally, we evaluated the relationship between individual APS and environmental variables using three models: linear, quadratic, and null, determining the best fit based on the lowest Akaike Information Criterion (AIC) value.

### 2.6 Loci annotation and Candidate Gene Identification

Loci identified as putative adaptative loci (see results section 3.5) were functionally annotated through Blast2GO v.6.0.3 (Conesa & Götz 2008). We compared the sequences adaptative loci were associated with genes that may be related to environmental tolerances (e.g., heat shock, immune system proteins, osmoregulation). We applied the BLASTx algorithm with a cut-off of E-value ≤1×10^−5^, homology of sequences of more than 75% and other parameters were used as default. Subsequent analysis only considered sequences with functional annotation (blasted, mapped and annotated).

## 3 Results

### 3.1 RAD sequencing and data filtering

After quality control, 2,359,848 (511,190 RADtag) putative loci were obtained for Stacks pipeline. Using the iterative filtering process (MAF and missing data), we identified a total of 2,328,560 (98.67%) putative singletons and 386 putative paralog loci; the latter were removed (1.24%). Further filtering by HWE discarded 13,215 loci (42.96%) and one SNP per tag (the highest F_ST_) discarded 5,582 SNPs (31.82%). A total of 11,961 SNPs (5.01%) were retained, from 202 individuals across the 10 sampling locations (Table 3).

**Table 3.**
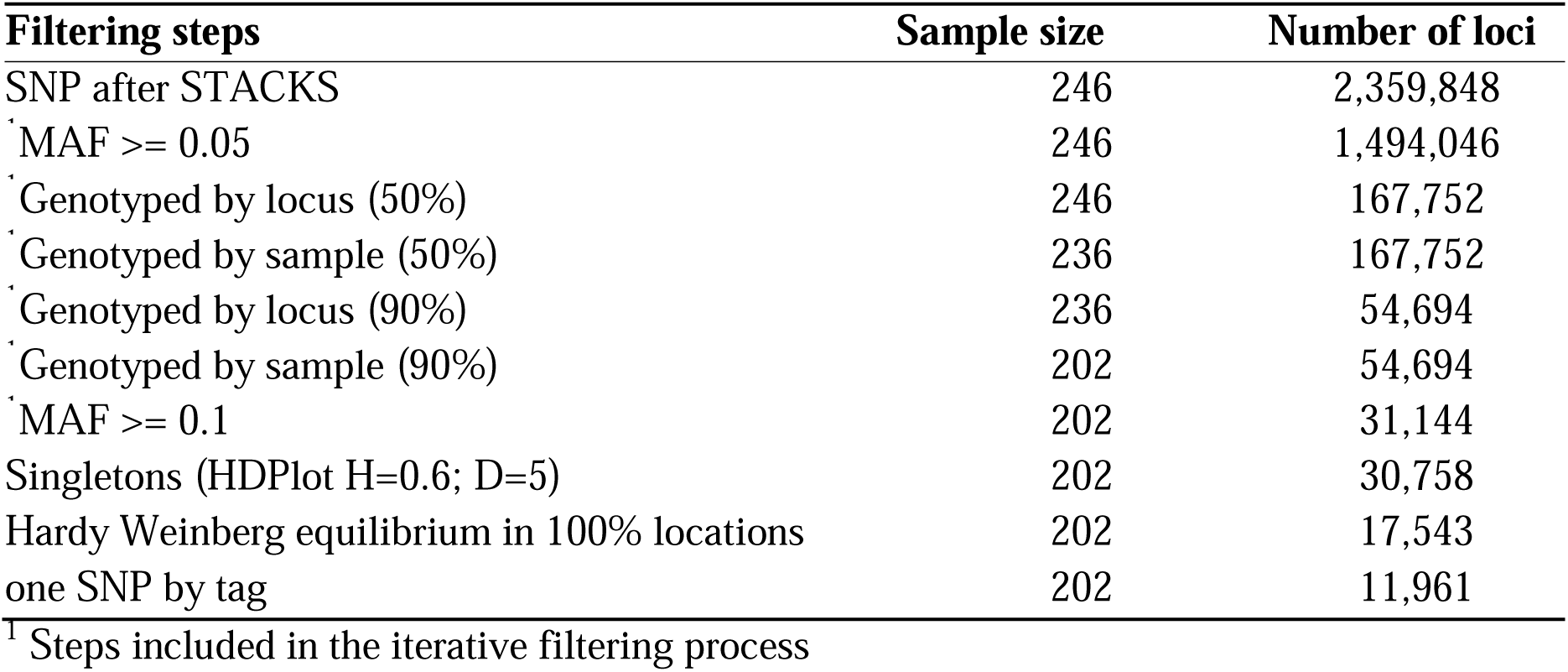
Description of steps used to filter SNPs.

### 3.2 Identification of loci putatively under selection

From population genetic differentiation (PGD) and genotype-environment association (GEA) we identified different putative adaptative loci. Out of the 11,961 loci, we identified 392 loci under positive selection by PGD, and 2,164 loci using the GEA approach. A total of 131 loci were identified as shared between the PGD and GEA approaches (Figure 2).

**Figure 2.**
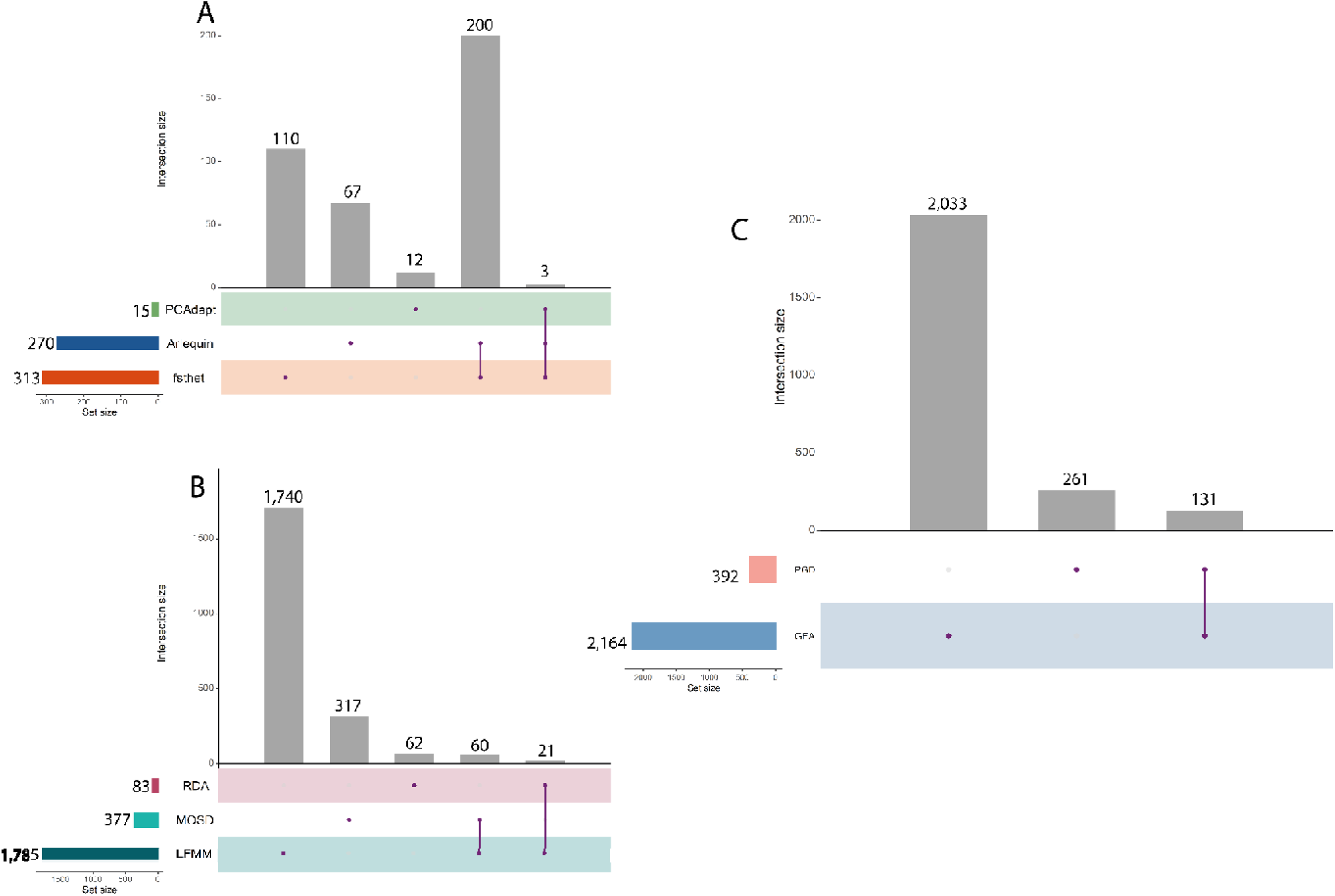
UpSet diagram: matrix layout for all intersections of software used. A) Loci detected by software based on Population Genetic Differentiation (PGD), B) Loci detected by software based on Environment Genotype-Association and C) Loci shared between PGD and GEA. The set size (horizontal bars) indicates the total number of loci detected by that software. The intersection size (vertical bars) indicates the number of loci per set

Within PGD approaches, *fsthet* detected 334 SNPs (2.79%) under balancing selection and 313 SNPs (2.62%) under divergent selection (Figure S2). *Arlequin* identified 270 SNPs under divergent selection (2.26%) (Figure S3). *PCAdapt* detected 15 SNPs (0.13%) under divergent selection (Figure S4). *BayeScan* and *OutFLANK* found no outlier. For GEA approaches, *LFMM* identified 1,785 loci under divergent selection, with the highest number of associations to nitrate concentration (n = 684 loci) (Figure S5). *MSOD* detected 377 loci, with the highest number of associations to temperature (n = 377 loci) (Figure S6). Finally, *RDA* detected 83 loci under divergent selection, with the highest number of associations to silicate (33 loci) (Figure S7). *Sam*β*ada* and *BayeScEnv* did not detect any correlation with environmental variables. Overall, across GEA approaches, the nitrate concentration (n = 844), temperature (n = 748), and salinity (n = 703) were the environmental variables with which most of the loci were correlated (Figure 3). Of the total of 2,164 that correlated with each of the six environmental variables, 783 of the loci were significantly associated with more than one environmental variable.

**Figure 3.**
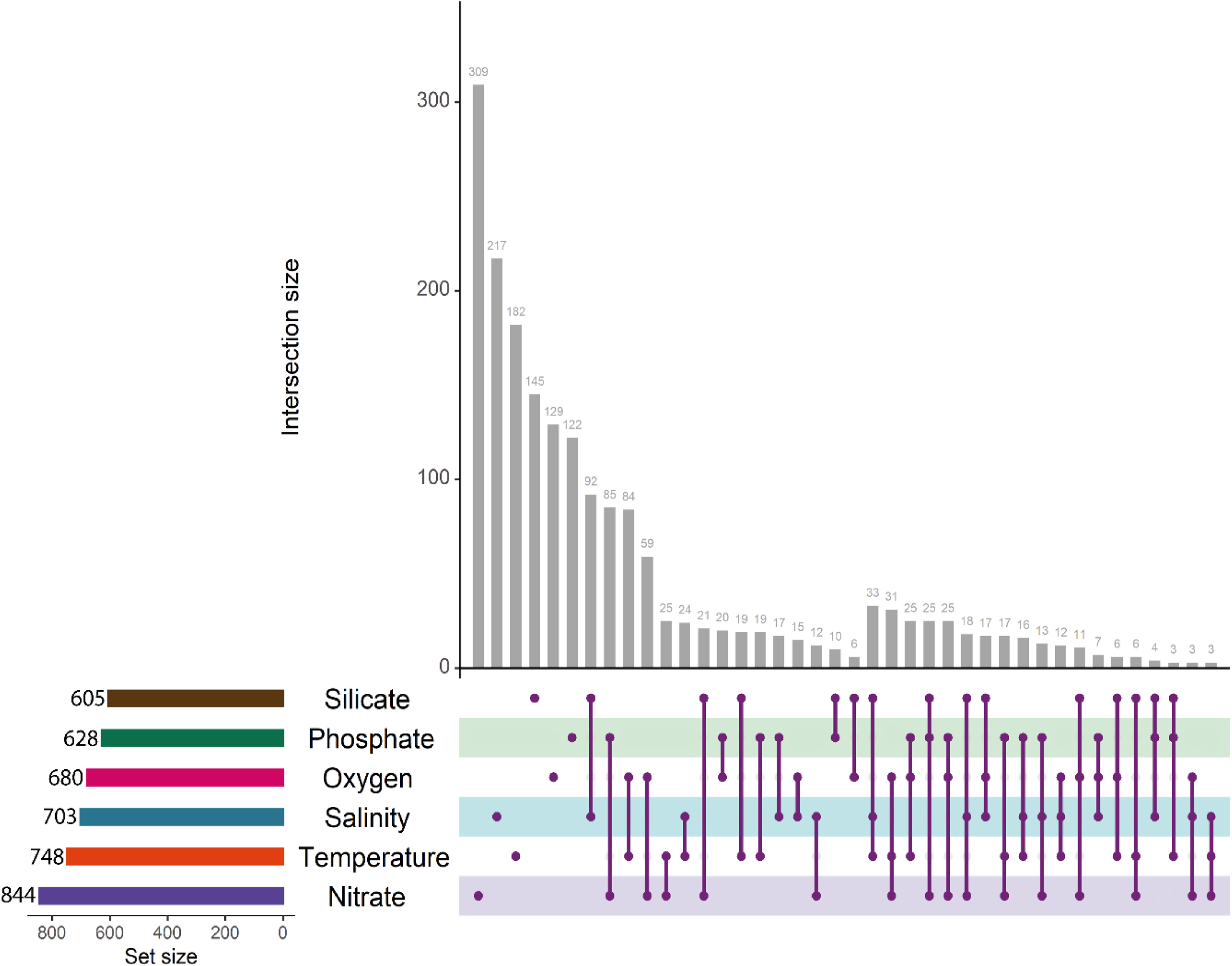
UpSet diagram: matrix layout for all intersections of environmental variables. The set size (horizontal bars) indicates the total number of loci correlated with that environmental variable. The intersection size (vertical bars) indicates the number of loci per set intersection, the purple dots represent the set of unique loci correlated only with that environmental variable and no other one, while the purple dots connected by a line represent loci shared by two, three, or more variables.

### 3.3 Assessment of genetic diversity and population structure

Summary statistics of genetic diversity revealed similar values for each location within each dataset (Figure S8). Most locations showed lower values of observed heterozygosity (H_O_) than expected heterozygosity (H_E_), and positive values of inbreeding coefficient F_IS._ For the adaptative loci dataset, H_O_ 0.3364 ± 0.0078 (C.I._95%_), H_E_ 0.3551 ± 0,0068(C.I._95%_), and F_IS_ 0.0528 ± 0.0120 (C.I._95%_). For the neutral dataset, H_O_ 0.3414 ± 0.0074 (C.I._95%_), H_E_ 0.3485 ± 0,0063 (C.I._95%_), and F_IS_ 0.0206 ± 0.0117 (C.I._95%_).

Pairwise-F_ST_ comparisons showed higher levels of genetic differentiation for adaptive loci (global F_ST_ = 0.0419 ± 0.0042 CI_95%_), than neutral loci (global F_ST_ = 0.0004 ± 0.0004 CI_95%)_. For adaptative loci, all pairwise differentiation values were highly significant (p-value ≤ 0.0004). While in neutral loci, 26 of 45 pairwise differentiation values showed significant p-values (Table 4).

**Table 4.** Pairwise F_ST_ (below diagonal) and p-value (above diagonal). Values between sampling site locations from North Patagonian fjord ecosystems in southern Chile using A) adaptative loci and B) neutral loci. Statistical significance was considered below 0.05 p-value.

| A) | REL | MAN | HOR | CHE | CHA | RMB | MEL | PUY | CIS | AYS |
| --- | --- | --- | --- | --- | --- | --- | --- | --- | --- | --- |
| REL | - | 0 | 0 | 0 | 0 | 0 | 0 | 0 | 0 | 0 |
| MAN | 0.0672 | - | 0.0004 | 0 | 0 | 0 | 0 | 0 | 0 | 0 |
| HOR | 0.0495 | 0.0126 | - | 0 | 0 | 0 | 0 | 0 | 0.0004 | 0 |
| CHE | 0.0653 | 0.0443 | 0.0493 | - | 0 | 0 | 0 | 0 | 0 | 0 |
| CHA | 0.0665 | 0.0502 | 0.0503 | 0.0531 | - | 0 | 0 | 0 | 0 | 0 |
| RMB | 0.044 | 0.0302 | 0.0227 | 0.0502 | 0.0502 | - | 0 | 0 | 0 | 0 |
| MEL | 0.0755 | 0.0343 | 0.0296 | 0.0521 | 0.0539 | 0.0313 | - | 0 | 0 | 0 |
| PUY | 0.0467 | 0.0403 | 0.0266 | 0.0546 | 0.0408 | 0.0286 | 0.033 | - | 0 | 0 |
| CIS | 0.0592 | 0.0338 | 0.0210 | 0.0546 | 0.0435 | 0.0248 | 0.0355 | 0.0234 | - | 0 |
| AYS | 0.0472 | 0.0442 | 0.0199 | 0.0553 | 0.0456 | 0.0269 | 0.0432 | 0.0263 | 0.0292 | - |

| B) | REL | MAN | HOR | CHE | CHA | RMB | MEL | PUY | CIS | AYS |
| --- | --- | --- | --- | --- | --- | --- | --- | --- | --- | --- |
| REL | - | 0 | 0 | 0.9997 | 0.7841 | 0.1541 | 0 | 0.0095 | 0.0763 | 0.0121 |
| MAN | 0.0018 | - | 0.0004 | 0.9909 | 0.3254 | 0 | 0 | 0 | 0.0008 | 0 |
| HOR | 0.0017 | 0.0011 | - | 0.998 | 0.9681 | 0.0018 | 0.0023 | 0 | 0.0174 | 0 |
| CHE | -0.0018 | -0.0011 | -0.0013 | - | 1 | 1 | 0.9814 | 0.9893 | 0.9994 | 0.9887 |
| CHA | -0.0394 | 0.0002 | -0.0789 | -0.0036 | - | 0.8519 | 0.8760 | 0.2556 | 0.9314 | 0.4809 |
| RMB | 0.0004 | 0.0016 | 0.0011 | -0.0022 | -0.0466 | - | 0.0003 | 0.0489 | 0.3379 | 0 |
| MEL | 0.0019 | 0.0019 | 0.0011 | -0.0011 | -0.524 | 0.0013 | - | 0 | 0.0070 | 0 |
| PUY | 0.0010 | 0.0020 | 0.0019 | -0.0011 | 0.0297 | 0.0006 | 0.0020 | - | 0.0004 | 0.0002 |
| CIS | 0.0006 | 0.0012 | 0.0007 | -0.0017 | -0.0684 | 0.0002 | 0.0010 | 0.0014 | - | 0.0006 |
| AYS | 0.0009 | 0.0017 | 0.0015 | -0.0010 | 0.2420 | 0.0012 | 0.0016 | 0.0012 | 0.0012 | - |

Evanno’s methods showed that the likely number of K clusters was K = 2 for adaptative loci (Figure S9) and K = 7 for neutral loci (Figure S10). For adaptative loci, a visual inspection revealed a clear separation of individuals for REL, MAN and MEL using K=2. We observed an additional cluster which appeared to correspond to individuals from CHE and CHA using K = 4 (Figure 4A). For neutral loci, *Structure* showed no clear geographical genetic structure (Figure 4B). Results from DAPC analyses revealed similar patterns to *Structure* (Figure 5). The adaptative dataset showed that the REL, CHA and CHE were separate from all other locations. The first two axis explain 44.9% of the total variation. For the neutral dataset, PET, CHA and CHE overlapped greatly with the other locations and much less structure overall was present, although MAN seemed to be less like the other nine locations. The first two axis explain 40.5% of the total variation (Figure 5).

**Figure 4.**
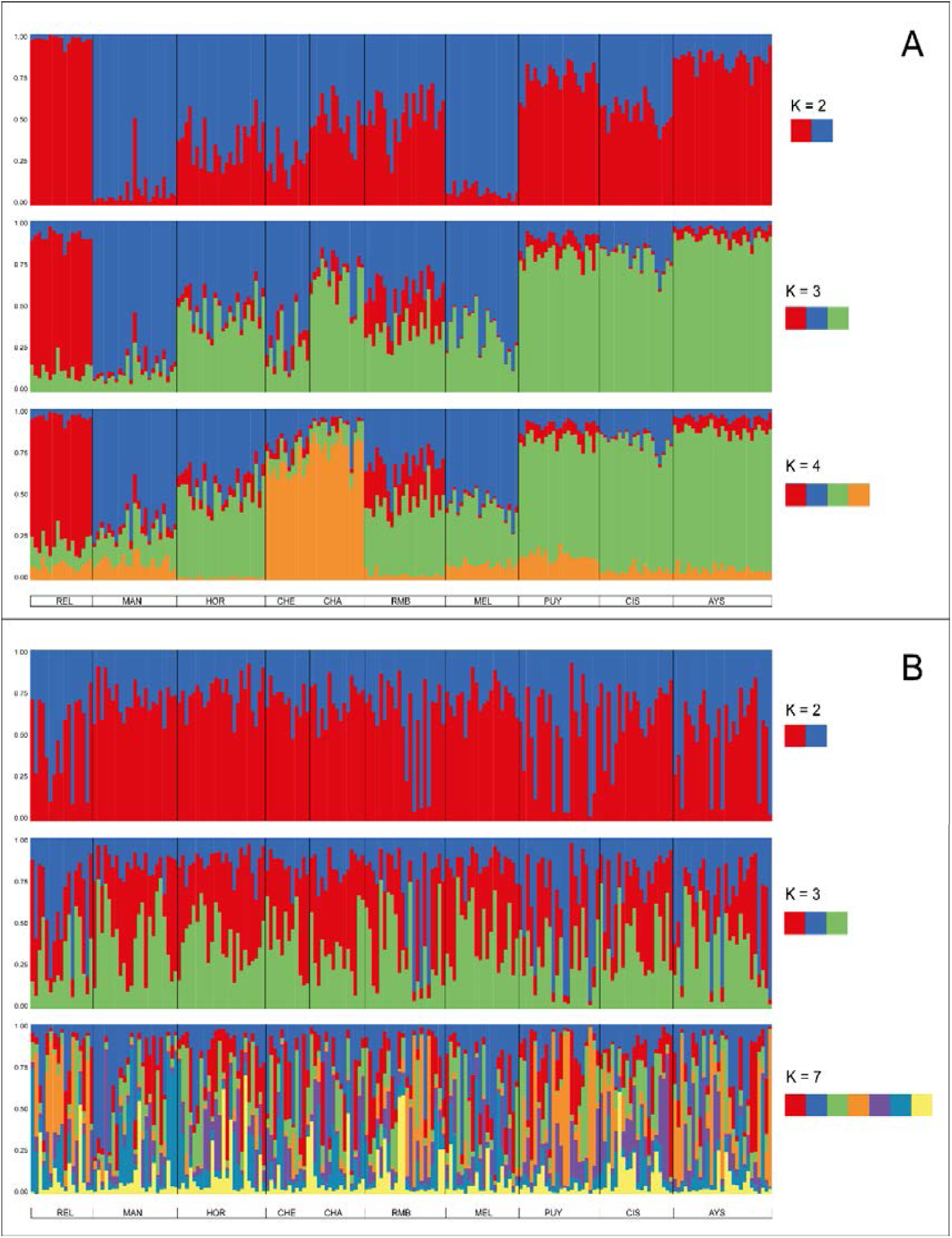
Results from clustering analysis of the *E. maclovinus.* Based on A) 131 putatively adaptative loci and B) 9,536 putatively neutral loci. Each vertical line represents an individual. The colors represent the proportion of inferred different locations. ΔK calculated for K ranging from 1 to 10, with each K repeated tenfold.

**Figure 5.**
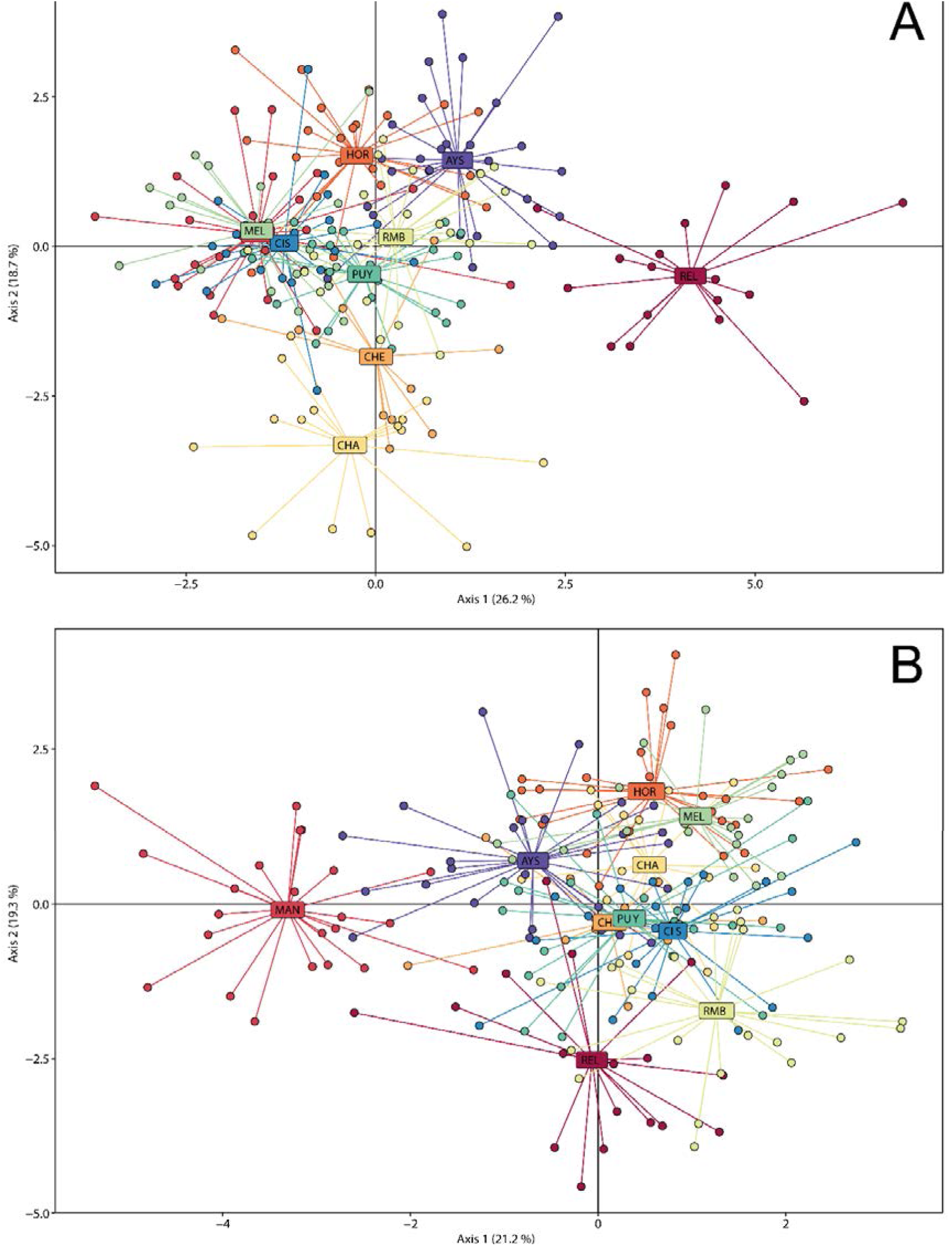
Scatterplot of discriminant analysis of principal component (DPCA). A) adaptative loci and B) neutral loci. Axes correspond to the first two discriminant 411 functions and the circles represent individuals colored-coded by sampling location.

The isolation by distance (IBD) analysis for adaptative significant isolation by distance loci revealed a moderate pattern (Mantel test: r = 0.3458, p-value = 0.0320, Figure 6A), where the scatterplot of local densities of distances showed only one consistent cloud of points, indicating a continuous cline of genetic differentiation. For neutral loci, we found no detectable relationship between geographic distance and genetic distance (Mantel test: r = −0.0435, p-value = 0.5760, Figure 6B).

**Figure 6.**
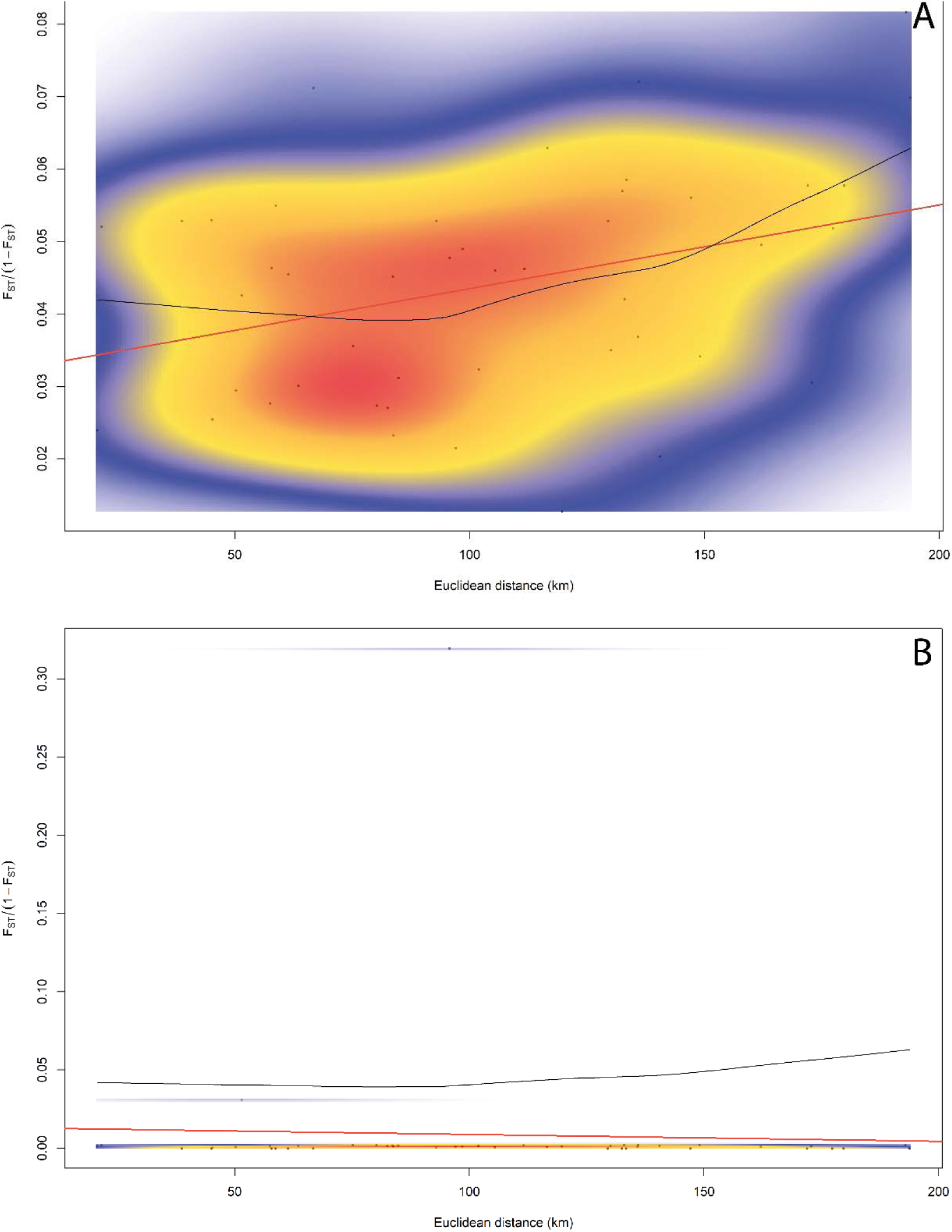
Isolation-by-distance IBD among locations. A) 131 adaptative loci and B) 9,536 neutral loci. Local density of points plotted using two-dimensional kernel density estimation. The black line is the trend estimated by Loess regression red line indicates the linear regression. Color gradient represents density of observations (low = blue, red = high).

### 3.4 Additive polygenic scores (APS)

The correlations between additive polygenic scores (APS) and environmental variables were all significant (p < 0.001). The R² values ranged from 0.29 for maximum temperature to 0.26 for both maximum salinity and minimum oxygen concentration. For all environmental variables, APS decreased with increasing temperature, salinity, and oxygen levels, respectively (Figure 7). The correlation between APS and environmental variables was best represented by linear models, except for maximum temperature, which fitted better with a quadratic model (Figure 7). The significance and R² values of the correlations calculated for mean and minimum temperature, salinity, and oxygen concentration yielded similar results, albeit with slight differences among subsets (Figure S11-S13).

**Figure 7.**
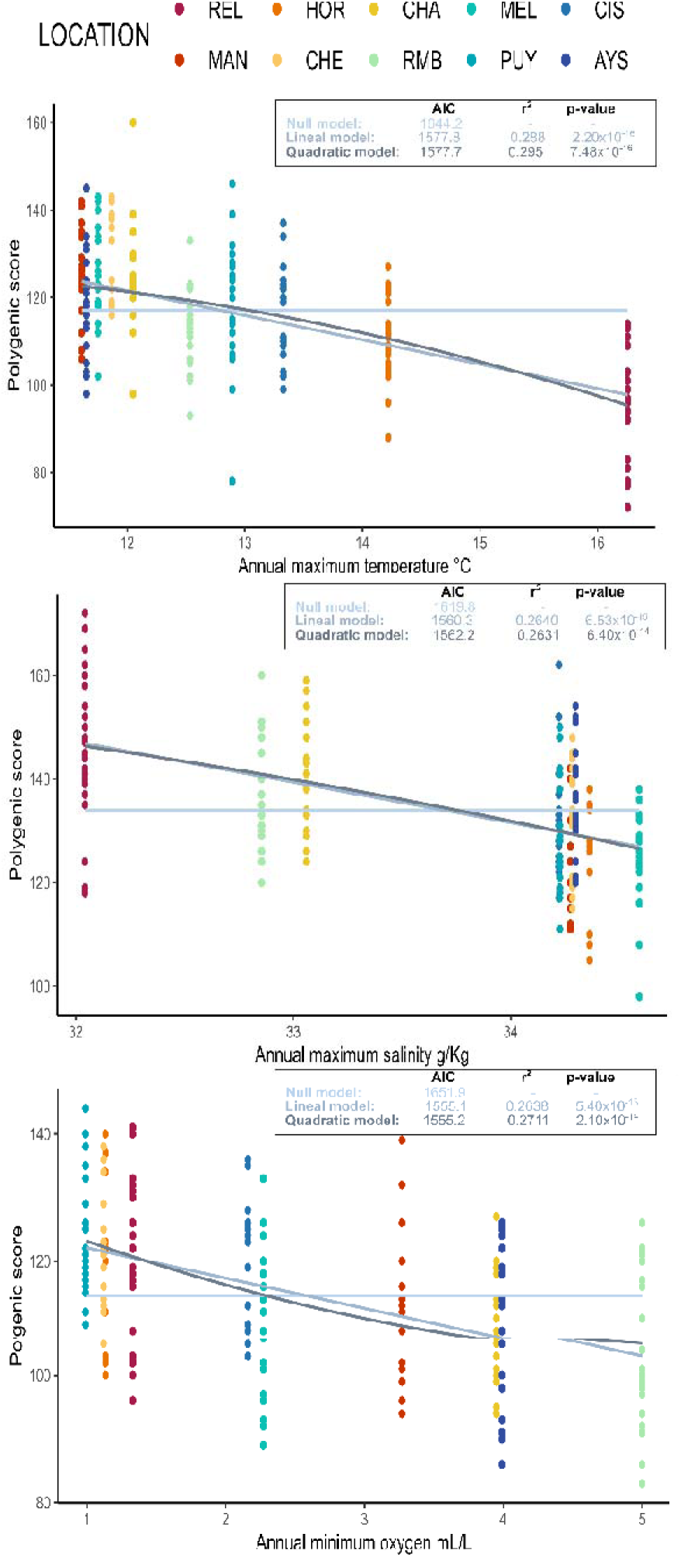
Correlations between additive polygenic scores (APS) based on A) Annual maximum temperature B) Annual maximum salinity and C) Annual minimum oxygen and 131 putative adaptative loci. Correlation coefficient (R^2^) and p-values and AIC are presented for each variable.

### 3.5 Gene ontology of candidate genes

The BLASTx analysis of the contigs containing the 131 adaptative loci shared by PGD and GEA against bony fish genomes resulted in significant hits (E-value ≤1×10^−5^) for 790 fish species. The BLASTx similarity results showed that 51 of the 131 contigs containing adaptative loci corresponded to known proteins in the non-redundant (NR) database (E-value 1e^−5^), of which 36 contigs were functionally annotated. The functional categorization of the annotated sequences involved biological process, cellular components, and molecular function (Figure 8).

**Figure 8.**
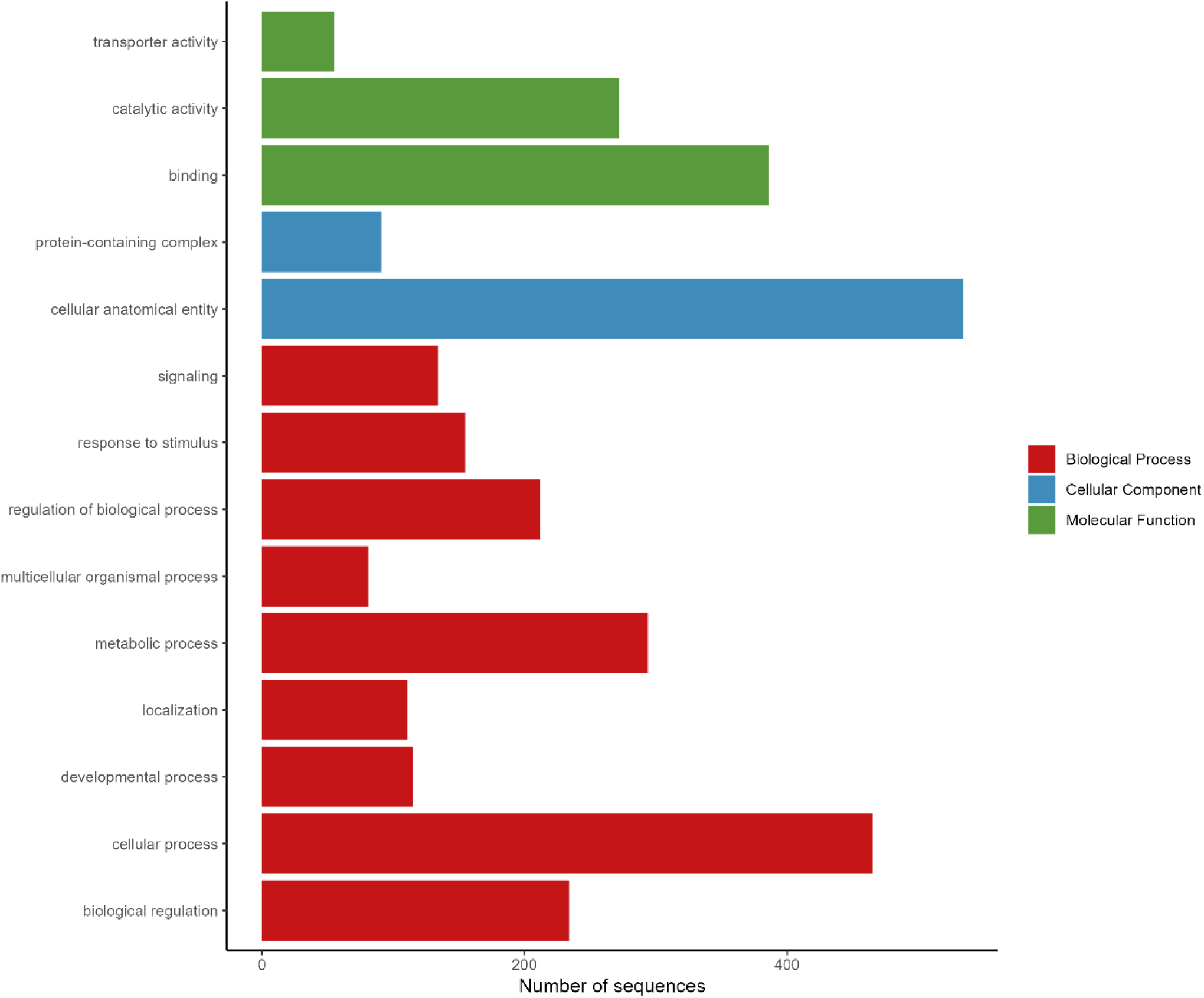
Putative functional categorization and distribution according to Blast2GO.

A second BLASTx analysis with all 2,425 putative loc i detected 512 contigs that were functionally annotated. Of these 512 sequences, three were already known to potentially play a role in the local adaptation (Table S1). The sequences of the *SNP 6291_214* and *44791_149* and belong to the solute carrier (SLC) family. SLC genes encode transmembrane transporters for inorganic ions, amino acids, neurotransmitters, sugars, purines and fatty acids, and other solute substrates (Dorwart *et al*. 2008). The SNP 42_59 is located in the vtgC (13489) gene, which is a member of the vitellogenins (Vtg) family. In egg-laying vertebrates such as fish, estrogens activate the hepatic synthesis of vitellogenin (Vtg) (Sumpter & Jobling 1995; Brian *et al*. 2008).

## 4 Discussion

This study provides comprehensive evidence for genomic signatures of local adaptation, of *Eleginops maclovinus* to heterogeneous environmental conditions in Northern Patagonia. We conducted a seascape genomics study using 11,961 RAD-sequencing markers and detected 2,164 putative adaptative loci through population genetic differentiation (PGD) and genotype-environment association (GEA) approaches. Using 131 loci shared by PGD and GEA, we observed that the putative adaptive loci collectively exhibit a highly significant association with various environmental variables, indicating polygenic selection. Additionally, we found contrasting patterns of genetic variation in neutral and adaptative loci that reinforce the possible local adaptation among high levels of gene flow. Subsequently, we detected one gene related to thermal adaptation and two genes that play a role in the osmoregulation. In the following, we describe how our study contributes to understanding the role of environmental heterogeneity as a driver of spatial patterns of putative adaptive genetic variation that could influence the local adaptation in *E. maclovinus*. Furthermore, we discuss methodological issues about software for detecting putative adaptative loci. Finally, we propose groups based on adaptative divergence for creating management units.

### 4.1 Evidence for spatially divergent selection

Evidence suggests thermal adaptation *E*. *maclovinus*, which juveniles exhibit significant eurythermal capacity, tolerating temperatures between 4 and 20°C (Vanella *et al*. 2012; Oyarzún *et al*. 2018, 2021). Short-term exposure to 4°C may reduce fitness (Vanella *et al*. 2012), while long-term exposure to 15-20°C elevates metabolism with tissue-specific modulation, without adverse physiological effects. Conversely, temperatures exceeding 25°C detrimentally impact physiology, growth, food intake, and energy substrates, mainly triglycerides (Oyarzún *et al*. 2021). These studies focus on juveniles; thus, temperature effects on early-stage adaptation remain uncertain for this species. Considering *E. maclovinus* spawns in spring (Brickle *et al*. 2005), higher temperatures could expedite maturation (e.g. Scott & Pankhurst 1992; Shimizu 2003; Hokanson 2011). Temperature affects reproductive processes’ thermal inhibition (Gillet *et al*. 1996; Tveiten *et al*. 2001; Okuzawa *et al*. 2003; Soria *et al*. 2008; Pankhurst & King 2010), including egg mortality (Van Der Kraak & Pankhurst 1997; Gagliano *et al*. 2007; Janhunen *et al*. 2010), embryonic development rate (Pauly & Pullin 1988; Rombough 1997; Smith *et al*. 2015), and larval fish metabolism, growth, and development (Houde 1989; Blaxter 1992; Benoît *et al*. 2000). Future research should examine temperature-induced shifts in selective mortality in early stages of *E. maclovinus*. Additionally, temperature may influence sexual inversion in protandrous hermaphrodites, as proposed by Baroiller et al. (1999). For instance, elevated temperatures increased gonadal aromatase levels, promoting female proportion in *Lates calcarifer,* another protandrous hermaphrodite (Athauda *et al*. 2012). This latter, although not identified in *E*. *maclovinus*, can be taken as a research question for future studies. The detection of potentially adaptive loci associated with temperature supports molecular adaptation via gene-related heterogeneity and gene duplication events, likely preserved in response to thermal fluctuations. Unlike most Antarctic notothenioids, *E. maclovinus* blood shows high hemoglobin (Hb) multiplicity, with one cathodal (Hb C) and two anodal components (Hb 1, Hb 2) (Coppola *et al*. 2010). This Hb diversity may facilitate oxygen transport adaptation to environmental changes and metabolic demands (Weber 1990; di Prisco & Tamburrini 1992; Feuerlein & Weber 1994; Weber *et al*. 2000; Fago *et al*. 2001). The maintenance of multiple Hbs in response to temperature differences in temperate waters, significantly larger than in Antarctica, is likely (Coppola *et al*. 2010; di Prisco *et al*. 2012). Moreover, the absence of antifreeze-glycoprotein (AFGP) coding sequence in *E. maclovinus* indicates no adaptation to near-freezing temperatures (Cheng *et al*. 2003).

Evidence suggests salinity acts as a selective agent driving genetic differentiation in *E. maclovinus* populations. Canales-Aguirre et al. (2022) identified associations between adaptive loci and environmental variables (salinity, oxygen, primary production, current velocity) in *E. maclovinus*. Juveniles are considered euryhalines, adapting to salinities between 5 and 45 PSU (Vargas-Chacoff *et al*. 2014, 2015a). An inverse relationship between the trypsin chymotrypsin−1 ratio (T/C ratio) and salinity was observed (Vargas-Chacoff *et al*. 2015b). The T/C ratio, linked to growth and nutritional status (Rungruangsak-Torrissen *et al*. 2006; Rungruangsak-Torrissen 2007), suggests better food utilization in low salinity, enhancing physiological energy. It is possible that food utilization and efficiency is better in *E. maclovinus* acclimated to low salinity, leading not to better growth, but to increased energy available for physiological processes such. Osmoregulation in low salinity consumes energy, affecting growth (Vargas-Chacoff *et al*. 2015b). Euryhaline fish adapt to varying osmotic challenges (Laverty & Skadhauge 2012). Climate change, habitat degradation, and anthropogenic activities increase salinity stress (Kültz 2015). Climatic models for Northern Patagonia predict reduced precipitation and increased temperature, affecting hydrological regimes and salinity (Aguayo *et al*. 2019). The salinity variability in microenvironments can have multiple impacts on reproduction, including DNA damage from salt stress or oxidative stress (Dowling & Simmons 2009), sperm motility (Beirão *et al*. 2018), incomplete egg binding (Herberg *et al*. 2018), disrupted egg activation (Ginsburg 1963) and egg penetration issues (Yanagimachi *et al*. 2017), and cause problems during DNA recombination (O, Wai-sum *et al*. 2006).

Our results indicate oxygen concentration exerts selective pressure on *E. maclovinus.* Environmental oxygen variability has been crucial in fish evolution, influencing anatomical, behavioral, and physiological adaptations for oxygen acquisition and delivery (Nikinmaa & Rees 2005). Fish have evolved to balance oxygen delivery while avoiding toxic reactive oxygen species (Taylor & McElwain 2010; Dong *et al*. 2011). They exhibit diverse hemoglobin structures and functions for oxygen transport (Powers 1974; Jensen & Weber 1985; Weber 1990; Jensen *et al*. 1998). Hemoglobin diversity in fish, including E. maclovinus, likely results from selection pressures like low oxygen levels(Nikinmaa & Rees 2005). Structural and functional analyses of *E. maclovinus* hemoglobin (Hb1Em) showed higher O_2_ affinity compared to *T. bernacchii,* indicating differences in temperature sensitivity and O_2_ release between sub-Antarctic and Antarctic notothenioids (Coppola *et al*. 2012). Silva & Vargas (2014) identified hypoxic zones in Northern Patagonia’s fjords (1.1–2.0 mL L-1); we emphasize the importance of studying the ability of to adapt to hypoxic conditions in *E. maclovinus* until now unclear.

In this study, we identified putative adaptive loci associated with the concentration of nitrate, phosphate, and silicate. Although there is no direct relationship between nutrient concentration and adaptive capacity in *E. maclovinus,* the nutrients could be a proxy for primary productivity. The productivity cycle of Patagonian channels and fjords has been divided into two seasons: a productive season characterized by a marked spring diatom bloom followed by pulsed productivity events dominated by a sequence of different phytoplankton species, and then a non-productive season that predominantly features small phytoplankton cells (Iriarte *et al*. 2007; Czypionka *et al*. 2011; Montero *et al*. 2011; Paredes & Montecino 2011). During the productive season and in the absence of persistent hydrodynamic phenomena that force nutrient replenishment in the euphotic zone, the supply of nitrate and phosphate to the well-lit surface layer would be the chief limiting factor for phytoplankton growth (Pizarro *et al*. 2000; Iriarte *et al*. 2007). The increased freshwater discharge into the Puyuhuapi channel during non-productive season coincides with increases in the concentrations of silicic acid and dissolved organic matter concentrations, as well as an increase in underwater light attenuation (Montero *et al*. 2017). A GEA analysis of Canales-Aguirre (2022) provided evidence for local adaptation associated to primary production on *E*. *maclovinus* likely connected to food availability (Gove *et al*. 2016; Fox *et al*. 2018). Similarly, multiple studies detected putative adaptive loci associated with chlorophyll (indicator of primary productivity), for example Boulanger *et al*. (2022) with *Mullus surmuletus,* Cayuela *et al*. (2020) with *Mallotus villosus,* Maselko et al. (2020) with *Sebastes alutus* and Diopere *et al*. with *Solea solea*.

### 4.2 Additive Polygenic Scores (APS)

For the Additive Polygenic Scores (APS) (Gagnaire & Gaggiotti 2016; Babin *et al*. 2017), we observed a strong correlation between polygenic scores and maximum temperature and salinity, and minimum oxygen. This suggests that *E. maclovinus* populations might be locally adapted to these environmental variables through a polygenic architecture. As we highlighted in the previous section (See 4.1.), we detailed the impact of these environmental variables on fish survival and development.

We found negative correlations between individual APS and all environmental variables, which could indicate directional selection, where different alleles are maintained in distinct environments with a relatively constant selection force along the environmental gradient. Specifically, the relationship between APS and maximum temperature was best represented by a quadratic model, suggesting stabilizing selection at intermediate temperatures. However, caution is advised with this conclusion as the difference in AIC between the linear and quadratic models was minimal (AIC _linear_ = 1577.8, AIC _quadratic_ = 1577.7).

The impact of temperature on additive genetic variation agreed with observations in the adaptive population structure membership plots, where the REL population in the northernmost latitudinal range with warmer temperatures separates from the other nine populations. For salinity and oxygen concentration, the linear model better explained the relationship with APS, indicating relatively constant selection along the salinity and oxygen gradients. These results are consistent with the observed adaptive population structure, where the CHE and CHA populations differentiate from the rest, with the CHE population exposed to higher oceanic salinities. Lastly, the minimum oxygen concentration is critical for fish reproduction and development, as hypoxia can delay embryonic development and hatching (Richards 2009; Wu 2009). However, it remains unknown how hypoxia affects the genetic expression and reproductive development of *E. maclovinus*.

While the APS approach has potential for evaluating the cumulative effect of potentially adaptive loci in complex environments, it has limitations. Without a reference genome, we cannot rule out the possibility that some potentially adaptive loci are linked. An annotated *E. maclovinus* genome would allow us to identify matches between adaptive loci and selected genomic regions, providing insights into specific genes involved in local adaptation (Manel *et al*. 2016). Additionally, comparing genotypic variation in potentially adaptive loci with individual phenotypic data is necessary to establish genotype, phenotype, and fitness associations (Barrett & Hoekstra 2011).

Our results suggest that maximum temperature, maximum salinity, and minimum oxygen concentration may drive spatially variable selection in *E. maclovinus* populations. Temperature, salinity, and oxygen concentration impact the environmental gradient experienced by eggs and larvae (Holliday 1969; Blaxter 1992; Brickle *et al*. 2005; Wu 2009; Lehtonen & Kvarnemo 2015), potentially influencing local adaptation. Future research should aim to detect molecular signatures of selection within a multilocus quantitative genetics framework to delve deeper into the genetic architecture of polygenic adaptive traits.

### 4.3 Finding functional genomics of candidate genes for local adaptation

Using BLASTx to identify genes linked to adaptation through a gene ontology approach (Haasl & Payseur 2016; Manel *et al*. 2016), we found a total of 2,425 potentially adaptative loci with 131 loci shared by PGD and GEA, only 512 and 36 loci were successfully annotated, respectively. In the 131 adaptative loci, only 39.69% of loci matched known sequences in the public database, highlighting the limitation of RAD sequencing in identifying candidate genes without a reference genome (Davey *et al*. 2013; Andrews *et al*. 2016; Catchen *et al*. 2017; McKinney *et al*. 2017a). Both adaptive datasets revealed loci linked to life-history traits, including thermal adaptation and osmoregulation.

Focusing on life history traits, we identified five key genes: *opa1, Stx16, FMVIA, foxj1b,* and *Lamc3*. The gene *opa1* regulates mitochondrial fusion and fission, preventing Cyc-c leakage and inducing apoptosis, which is linked to kidney injury (Song *et al*. 2017; Yingjie *et al*. 2019; Herkenne *et al*. 2020; Wang *et al*. 2020; Cui *et al*. 2021; Li *et al*. 2022). The *Stx16* is involved in the development of inducible defense structures in *Daphnia pulex* under predation stress (Spanier *et al*. 2010). High *FMVIA* expression in the retina of striped bass suggests a role in retinal motility (Breckler *et al*. 2000). The *foxj1b* gene is essential for normal laterality in zebrafish embryos, affecting heart and organ positioning (Tian *et al*. 2009). Finally, *Lamc3* plays a role in the development of the parachordal chain in zebrafish (Eve & Smith 2017).

Expanding our search to adaptive loci, we identified three key genes: *vtg3* (linked to temperature adaptation) and *slc39a6, slc8a2a* (associated with salinity adaptation). *Vtgs* are specialized large lipid transfer proteins produced by the liver and transported to the ovary (Babin *et al*. 2007). The *Vtg* family includes type-I (*Vtg 1, 4, 5, 6,* and *7*), type-II (*vtg2* and *vtg8*), and type-III (*vtg3*) (Yilmaz *et al*. 2018). Despite being a minor and divergent form, *vtg3* significantly contributes to zygote and early embryo development (Yilmaz *et al*. 2019). Increased temperature elevates estrogen-induced *vtg* expression in juvenile *Salmo trutta* (Körner *et al*. 2008), suggesting a similar role for *vtg3* in *E. maclovinus* related to thermal adaptation. The gene *slc39a6 (LIV-1)* maintains zinc homeostasis, essential for cellular growth and differentiation (Taylor & Nicholson 2003). Low salinity increases *slc39a6* expression in tiger puffer (*Takifugu rubripes*) (Jiang *et al*. 2020), and intracellular zinc homeostasis controls T-cell survival in zebrafish by inhibiting apoptosis (Zhao *et al*. 2019). The *SLC8* gene family (*NCX*) maintains cellular Ca2+ homeostasis (Khananshvili 2013). *Slc8a2a*, an isoform identified in zebrafish, is key for Ca2+ excretion by seawater fish kidneys, with upregulated transcription during seawater acclimation (Liao *et al*. 2007; Islam *et al*. 2011). These genes indicate positive selection associated with salinity in *E. maclovinus* in Northern Patagonia.

We did not identify genes whose functions are supposed to be affected by the concentration of oxygen or nutrients (nitrate, phosphate and silicate). However, many outlier sequences did not have BLASTx results and may represent important proteins that have not been adequately annotated in fish to date.

### 4.4 Comparing approaches for detecting gen omic signal of selection

The outcomes from the ten programs varied widely in terms of the number and identity of adaptative loci detected in this study. Such variations among methods were expected since their algorithms and prior assumptions considerably differ. Combining population genetic differentiation (PGD) and genotype-environment association (GEA) approaches to detect adaptative loci of local adaptation not only reduces false-positive discoveries, but also maximizes our chances of detecting potential signals of selection (François *et al*. 2016). Overall, we found that GEA approach identified more adaptative loci (2,164 loci) than PGD approach (392 loci). These outcomes are in agreement with a simulation study demonstrating that the GEA approach has more power to detect loci under divergent selection than PGD approach (de Villemereuil *et al*. 2014), which is not surprising given that GEA uses multiple environmental variables (six variables in our case) to depict signals of selection. This trend has been reported in multiple studies focused on detecting signs of local adaptation in marine populations (Benestan *et al*. 2016; Flanagan *et al*. 2016; Dalongeville *et al*. 2018; Bekkevold *et al*. 2020; Cayuela *et al*. 2020; Segovia *et al*. 2020).

Inside of PGD approach, *fsthet* detected the largest number on putative adaptative loci (n= 313 loci). This software generates smoothed quantiles for the F_ST_–heterozygosity relationship from the empirical distribution (Flanagan & Jones 2017). Using an empirical distribution itself has the advantage of not assuming a particular distribution or model of evolution. For example, *FDIST2* and *LOSITAN* used an idealized island model, which would not be appropriate considering the geography of Northern _Patagonia. Besides, this_ feature is appealing because the F_ST_–heterozygosity distribution changes based on many potentially unknowable features of population structure (Flanagan & Jones 2017). Though, *fsthet* decreased the average proportion of false positives compared with *LOSITAN*, we did not find any studies that compared the effectiveness of *fsthet* with respect to other software that considers the hierarchical population structure, such as *Arlequin*. The *fsthet* program is not recommended when migration rates are low because the neutral loci saturate the full distribution of potential F_ST_–heterozygosity values (Flanagan & Jones 2017). In our case, the relative migration rates showed high gene flow between locations of *E. maclovinus* (Canales-Aguirre *et al*. 2022). The *Arlequin* identified 270 loci (200 shared with *fsthet*). However, results of a simulation study showing that *Arlequin* consistently found more outliers and had the highest type I and type II errors in their simulation scenarios in compared with other methods such as *BayeScan*. Simulated data suggested that *Arlequin* does not perform well in situations where adaptive variation contrasts with patterns of neutral variation, likely due to the effect of hierarchical grouping based on neutral markers (Narum & Hess 2011). The *PCAdapt* was the more conservative (n= 15 loci), whereas *BayeScan* and *OutFLANK* did not detect any outliers. Population structure and the presence of admixture individuals affect the power of identifying adaptive loci with *Bayescan* more than *PCAdapt* (Luu *et al*. 2017). This latter, agreed with our data where we found admixture individuals in the whole SNPs database. The presence of mixed individuals is supported by the high relative migration rates estimated by Canales-Aguirre *et al*. (2022). Additionally, *BayeScan* could decrease power with few populations (n <30 Aguirre-Liguori *et al.* 2020) and weak selection, when populations 694 deviate from the island model of migration and/or in cases of smaller sample size (Foll & Gaggiotti 2008; Narum & Hess 2011).

Inside the GEA approach, *LFMM* detected the largest number of potentially adaptive loci (n= 1,785 loci). Considering that *LFMM* identified 14.92% of the loci as adaptive, it is likely that we will find false positives among them. The main reason is that geographic and demographic processes can lead to patterns that mimic those observed due to selection. In fact, de Villemereuil *et al*. (2014) found high rates of false discovery in some scenarios with complex, hierarchical structures and polygenic selection; although showed that *LFMM* provides the best compromise between detection power and error rates in situations with complex hierarchical neutral genetic structure and polygenic selection. However, Frichot et al. (2013) found that *LFMM* has low rates of false positives and negatives and that it performs slightly better than *Bayenv* in detecting weak selection. Finally, Lotterhos & Whitlock (2015) showed that *LFMM* is quite robust to various sampling designs and underlying demographic models. The *MSOD* detected considerably fewer potentially adaptive loci (n=377 loci). However, the main advantage of this program is that using Moran spectral randomization (MSR) for significance testing, avoids the inflation of type I errors due to spatial autocorrelation, whether resulting from gene flow or the spatial structure of the environmental factors that induce selection (Wagner & Dray 2015; Wagner *et al*. 2017). Moreover, spatial outlier detection worked best for scenarios of IBD, even in the presence of large-scale spatial genetic structure (Wagner *et al*. 2017) as in this study.

#### LFMM MOSD

The main difference is the multivariate approach of *RDA*, while *LFMM* and *MSOD* are univariate, meaning that they test one genetic marker at a time, and may also test only one environmental predictor at a time (Rellstab *et al*. 2015); *RDA* identifying covarying loci associated with the multivariate environmental variables (Legendre & Legendre 2012). The decrease in the number of loci detected by *RDA* is because it restricts the number of environmental variables tested based on the variance inflation factor. In the case of univariate programs, we tested six environmental variables with four statistics for each but restricted to one statistic per environmental variable for *RDA*, limiting the number of possible correlations. However, simulated data showed that *RDA* has higher true-positive and lower false-positive rates compared to *PCAdapt* and *LFMM* (Forester *et al*. 2018; Capblancq *et al*. 2018). Furthermore, *RDA* has one important limitation: it models linear relationships among response and predictors, meaning that nonlinear relationships will not be detected (Capblancq & Forester 2021). Finally, we did not detect any putative adaptive loci using the *BayeScEnv* and *Sam*β*ada* programs. Considering that *BayeScEnv* produces fewer candidate loci than *BayeScan* (de Villemereuil & Gaggiotti 2015), it is to be expected that if *BayeScan* did not detect any loci *BayeScEnv* either. To date, we have not identified any studies that compare the statistical power or control of FDR of *BayeScEnv* with respect to other programs. Therefore, we cannot conclude whether *BayeScEnv* is more or less sensitive to certain characteristics such as hierarchical population structure, isolation by distance, or strength of selection. Due to the recentness of the *Sam*β*ada*, we did not identify comparative studies that allow us to evaluate the advantages and disadvantages of this with respect to other GEA approaches.

Continued testing and comparing PGD and GEA approaches is needed in simulation frameworks that include more population structure, multiple selection surfaces and genetic architectures that are more complex than the multilocus selection response modelled here. Overall, we highlight the importance of combining PGD and GEA approaches that consider multiple scenarios of adaptive variation, to diminish type I and II errors and avoid conclude erroneously.

### 4.5 Implications for management and conservation

The 11,961 filtered loci genotyped for 202 individuals (the largest population genomic data set reported so far for *Eleginops maclovinus*) offers high resolution for creating management policies and decisions, with respect to patterns of adaptive genetic variation. Although neutral loci did not reveal a clear genetic structure among locations, analysis of 131 shared adaptive loci indicated genetic differentiation separating the populations into four groups: REL, CHE, CHA, and a fourth group comprising the remaining seven sampling locations. Note that all neutral pairwise F_ST_ values were close to zero, similar to showed by Canales-Aguirre *et al*. (2022) and also from fish and other marine species (Nayfa & Zenger 2016; Asaduzzaman *et al*. 2019b; a; O’Leary *et al*. 2021). The interpretation of extremely small F_ST_ is not straightforward (Waples & Gaggiotti 2006; Conover *et al*. 2006; Waples *et al*. 2008), meaning it is unclear what the implications of the patterns of weak neutral differentiation may be for demographic connectivity of subpopulations. Nevertheless, the signals of putatively adaptive genetic differentiation between the ten locations were significant. Our study demonstrates how the environment shapes the distribution of adaptive genetic variation across space for *E*. *maclovinus*; showing that REL, CHE and CHA retain adaptive characteristics different from the rest. This can have important implications for prioritizing sites for protection, where the level of adaptive genetic variation could be an indicator of the evolutionary resilience of populations (Bonin *et al*. 2007; Sgrò *et al*. 2011). Moreover, understanding the spatial distribution of putatively adaptive genetic variation can inform the selection of specific sites for protection within marine reserves to maintain the adaptive potential and evolutionary resilience of wild populations faced with environmental change (von der Heyden 2017). The adaptive divergence reported here appears to be underpinned by a variation in seawater temperature and oxygen concentrations in the Northern Patagonia (Iriarte *et al*. 2014; Torres *et al*. 2014; Pérez-Santos *et al*. 2014; Silva & Vargas 2014) and complex patterns of population structure and putatively adaptive diversity among *E. maclovinus* locations that we showed here are not recognized in current *E. maclovinus* management units. To date, there is no management plan for *E. maclovinus* in Chile, only regulations for the type of fishing gear used. Future research could use neutral and adaptative datasets to determine management units using adaptative (e.g., population adaptative index, adaptative score) and neutral (e.g., expected heterozygosity, local differentiation) metrics as suggested by Xuereb et al. (2021). In the Chilean Northern Patagonia, temperature and heat content are predicted to increase and freshwater river inputs are predicted to decrease due to climate change (Aguayo *et al*. 2019). As such, the spatial patterns of genetic variability observed in this study can be information conservation planning decisions by identifying for protection populations containing higher levels of segregating polymorphisms associated with environmental conditions (e.g., temperature and salinity) that are prone to change in the future.

## Data Archiving Statement

Genotypes are available in https://github.com/Canales-AguirreCB/Claure_AdaptiveGenetics_MS

## Footnotes

1 Conservative temperature is a thermodynamic property of seawater. It is derived from the potential enthalpy and is recommended under the TEOS-10 standard (Thermodynamic Equation of Seawater-2010) as a replacement for potential temperature as it more accurately represents the heat content in the ocean.

2 The Absolute Salinity is the mass fraction of dissolved non H_2_O material in a seawater sample at its temperature and pressure and is expressed in g Kg^−1^

