## supplementary material for "Genomic signals of local adaptation in *Eleginops maclovinus* from Northern Chilean Patagonia"

**Genomic signals of local adaptation associated with environmental variables in *Eleginops maclovinus* from Northern Chilean** **Patagonia**

C. Eliza Claure ^1,2,3,4^, Wesley A. Larson ^5^, Garrett D. McKinney ^6^, J. Dellis Rocha ^7^, José M. Yáñez^8^, Cristian B. Canales-Aguirre ^1,2^

^1^ Centro i~mar, Universidad de Los Lagos, Camino a Chinquihue 6 km, Puerto Montt, Chile

^2^ Núcleo Milenio INVASAL, Concepción, Chile

^3^ Programa de Magister en Ciencias, mención Producción, Manejo y Conservación de Recursos Naturales, Universidad de Los Lagos

^4^ Programa de Doctorado en Ciencias, mención Conservación y Manejo de Recursos Naturales, Universidad de Los Lagos

^5^ National Oceanographic and Atmospheric Administration, National Marine Fisheries Service, Alaska Fisheries Science Center, Auke Bay Laboratories, Juneau, Alaska, USA

^6^ Washington Department of Fish and Wildlife, Seattle, Washington, USA

^7^ Escuela de Obstetricia, Facultad de Ciencias para el Cuidado de la Salud, Universidad San Sebastián Sede de La Patagonia, Puerto Montt, Región de Los Lagos

^8^ Facultad de Ciencias Veterinarias y Pecuarias, Universidad de Chile, Av Santa Rosa 11735, La Pintana, Santiago 8820808, Chile

*Corresponding author:

Cristian B. Canales-Aguirre

Centro i~mar, Universidad de Los Lagos

Camino a Chinquihue 6 km, Puerto Montt, Chile

### Supplementary data

#### Supplementary Figures

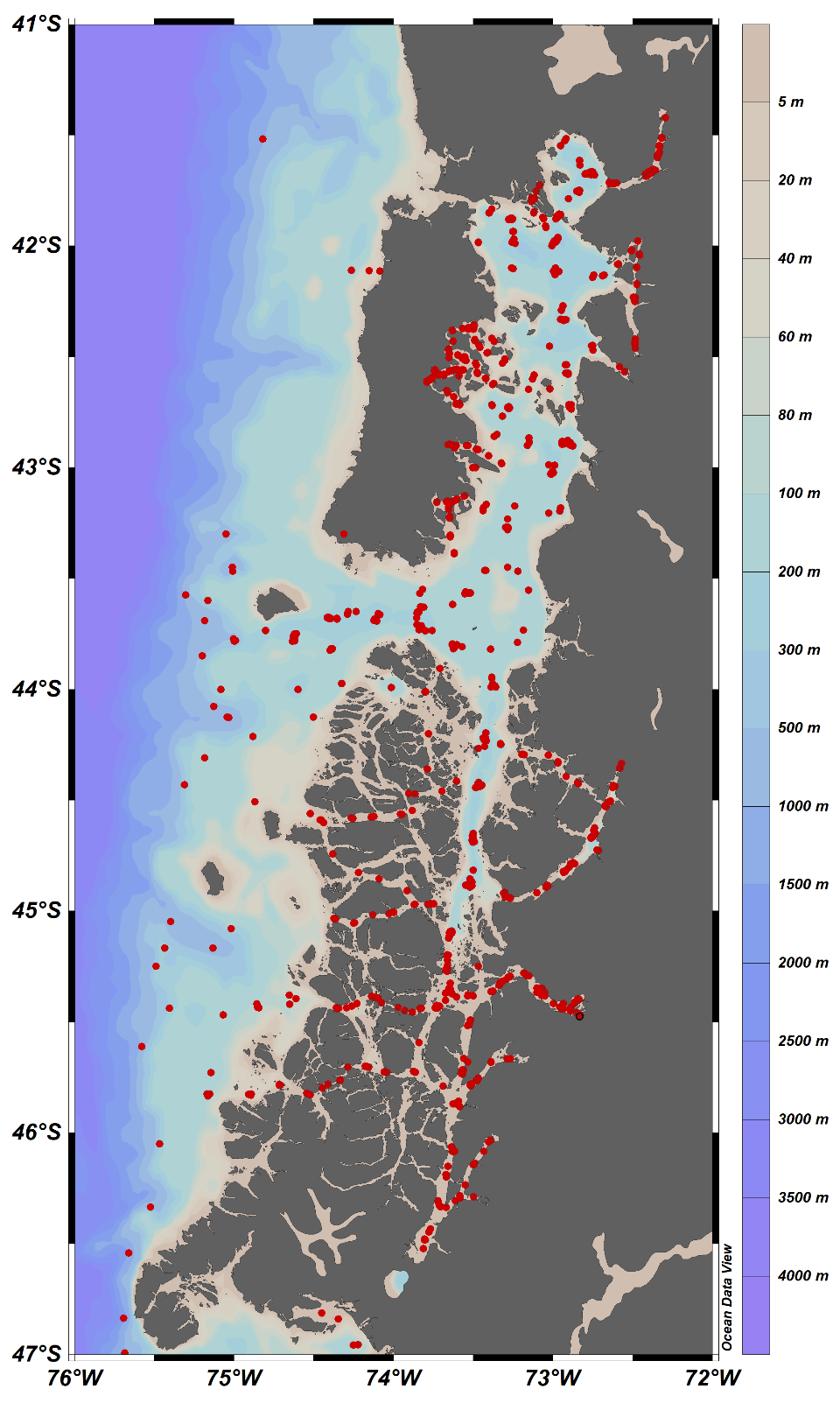

Figure S 1 Distribution of oceanographic station from CIMAR campaigns.Map showing the distribution of stations sampled during oceanographic campaigns CIMAR 1 until CIMAR 24 carried out in Northern Patagonia fjords and channels from.

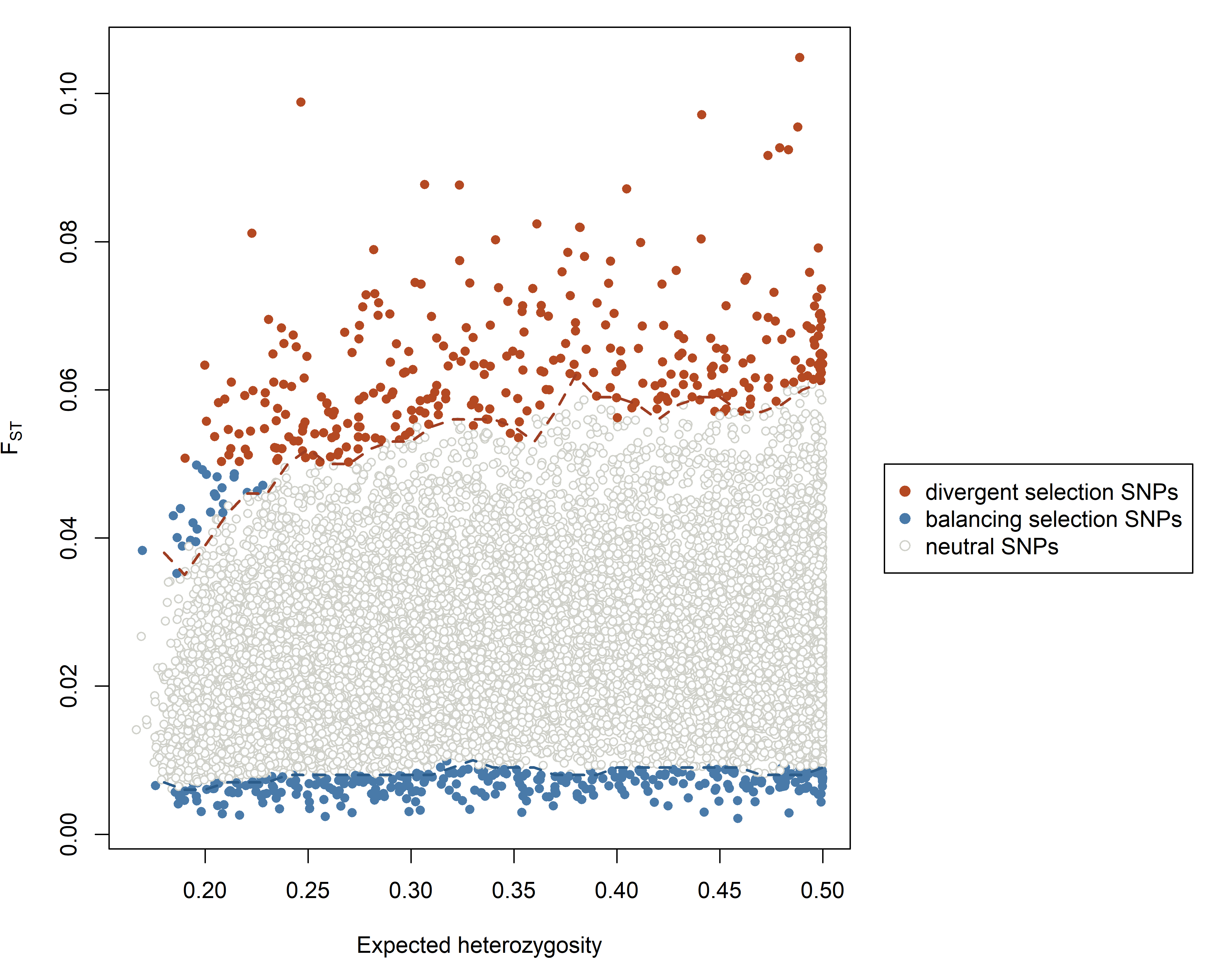

Figure S 2 Graph of the distribution of expected heterozygosity values in relation to F_ST_ values. Orange, blue and gray points represent putatively under divergent selection, under balancing selection, and neutral loci respectively using a threshold of 0.05. The lines show 95% smoothed quantiles calculated by *fsthet*.

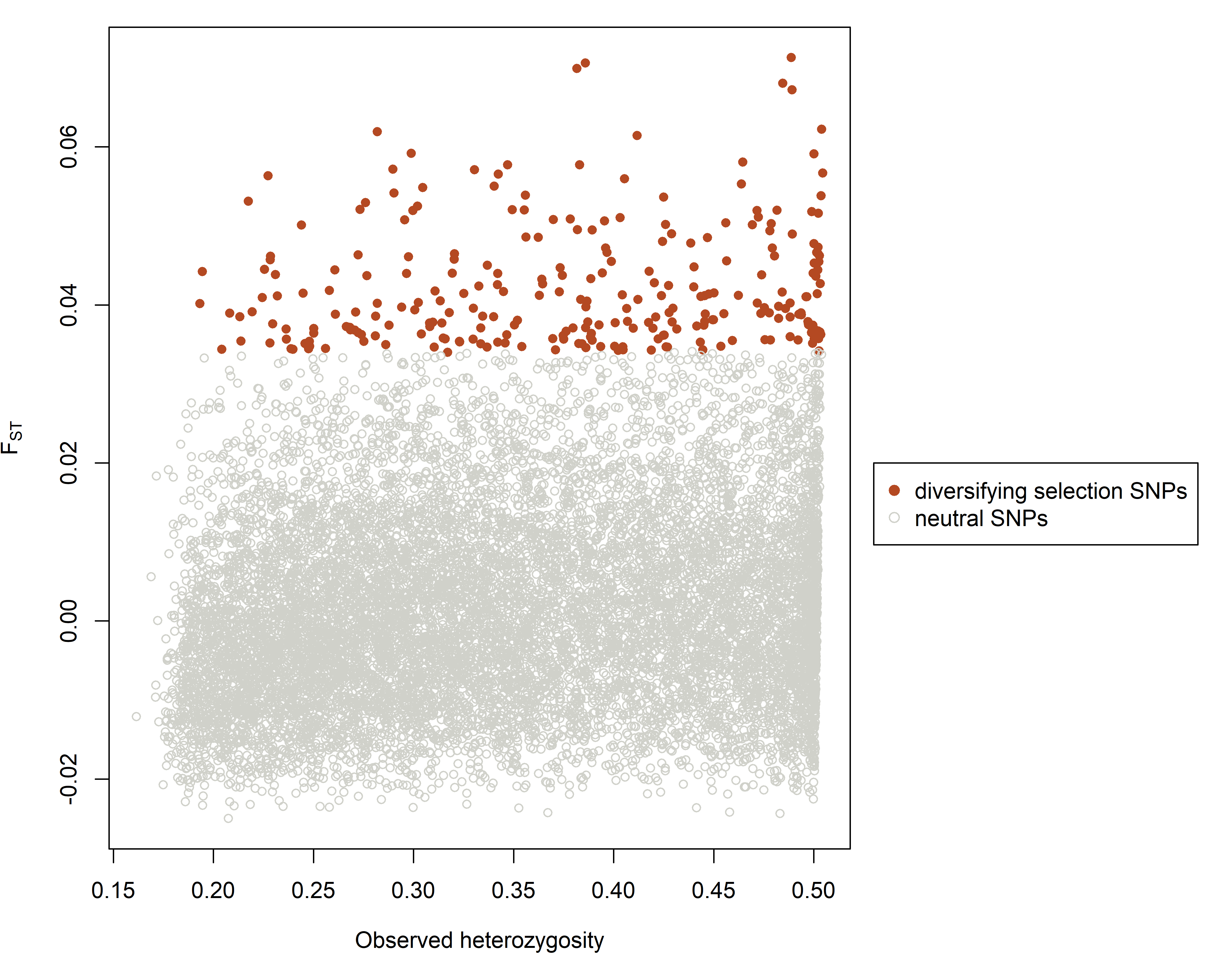

Figure S 3 Outlier loci under the hierarchical structure.

Model using *Arlequin 3.5.2.2.* are above the 95% quantile: F_ST_ and observed heterozygosity, putatively neutral loci are plotted as gray circles and putative loci under divergent selection are represented as orange circles. The putative loci under balancing selection were not identified by this software.

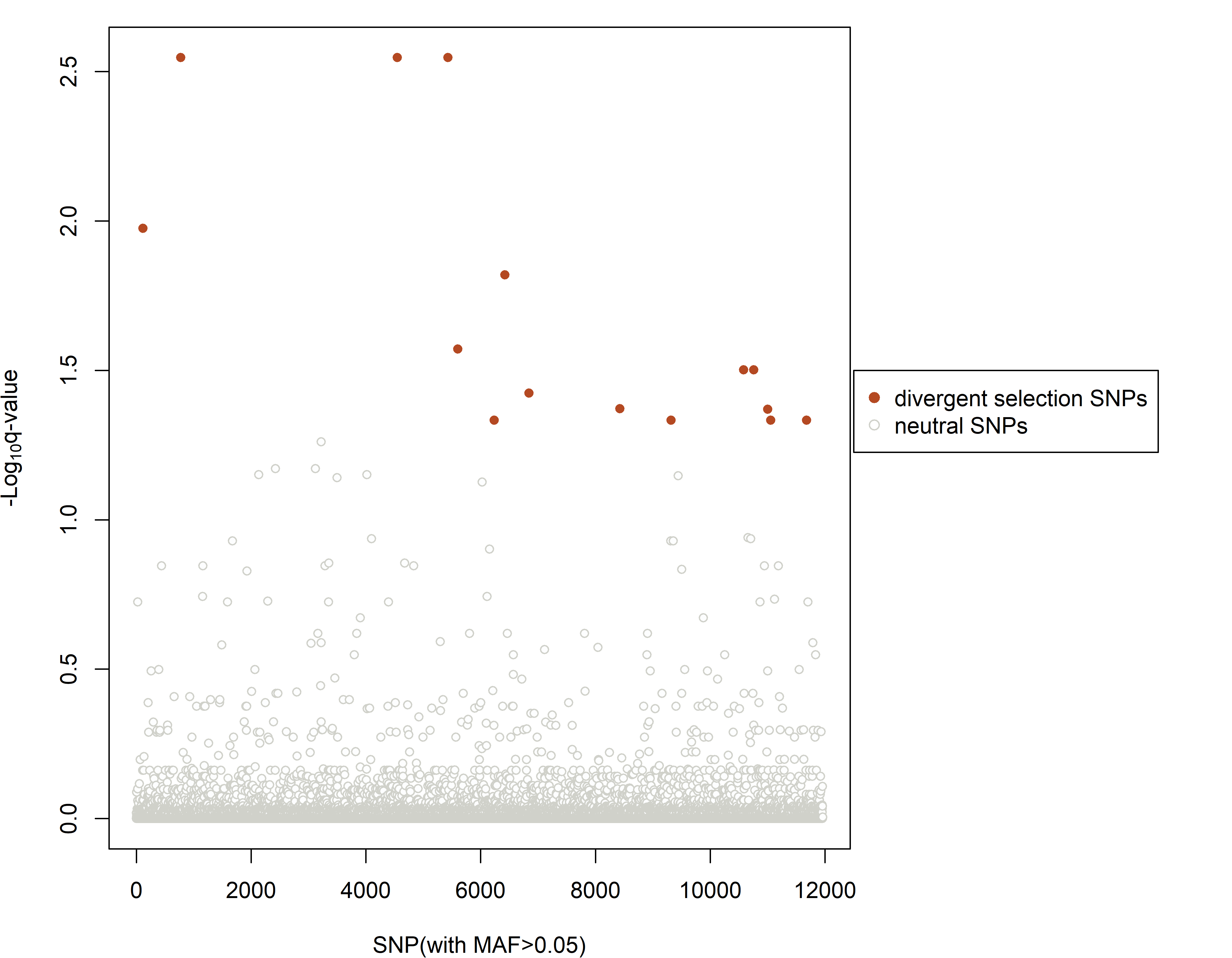

Figure S 4 Manhattan plot showing the distribution of putative adaptative loci.

Those obtained from *PCAdapt* where Y-axis represents p-values and putative adaptative loci above α = 0.05 are represented as orange circles.

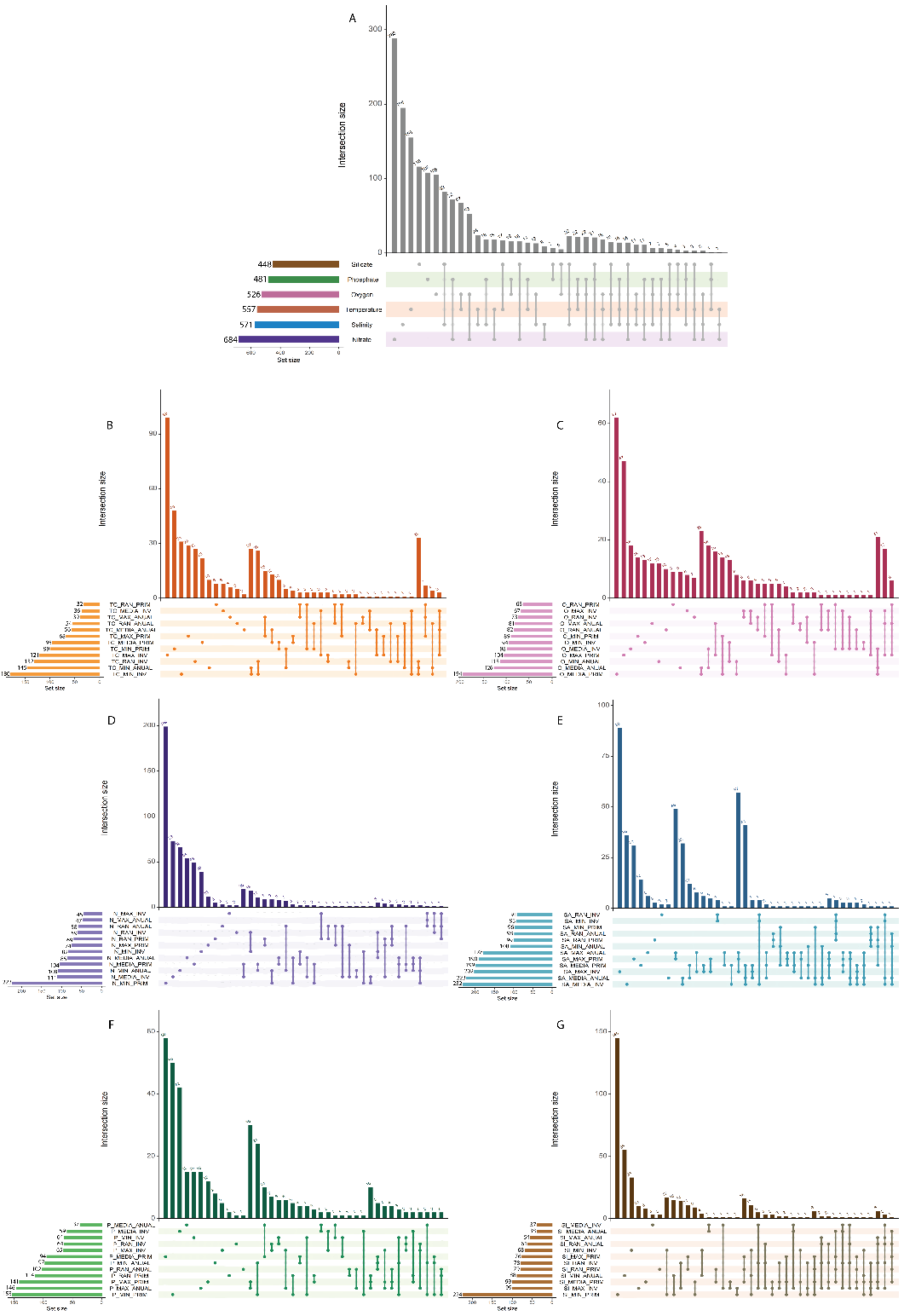

Figure S 5 UpSet diagrams for all intersections of adaptative loci detected by *LFMM*.

A) Summary of all putative adaptative loci. Disaggregated of results by B) Conservative temperature (ºC), C) Oxygen concentration (mL/L), D) Nitrate concentration (µM), E) Absolute salinity (g/Kg), F) Phosphate concentration (µM) and G) Silicate concentration (µM). The set size (horizontal bars) indicates the total number of loci correlated with that environmental variable. The intersection size (vertical bars) indicates the number of loci per set intersection, the dots represent the set of unique loci correlated only with that environmental variable and no other one, while the dots connected by line represent loci shared by two, three, or more environmental variable.

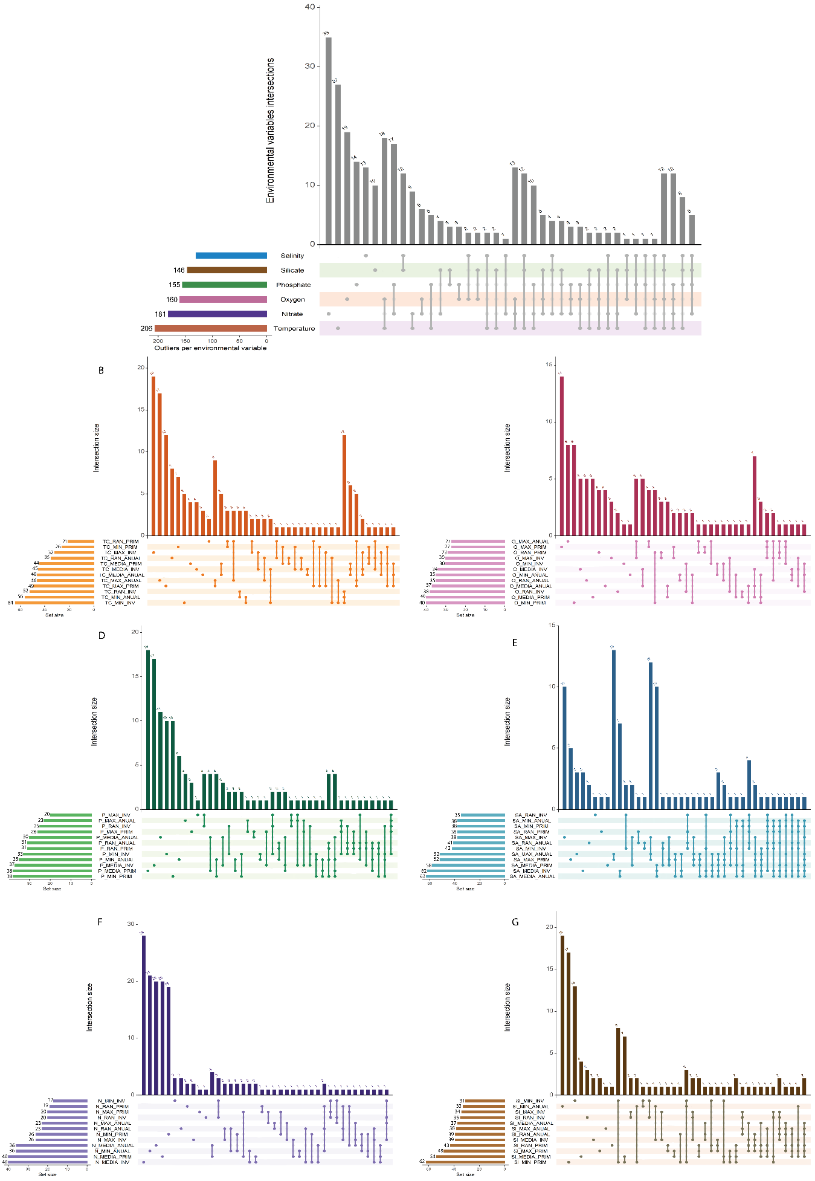

Figure S 6 UpSet diagrams for all intersections of adaptative loci detected by *MSOD.*

A) Summary of all putative adaptative loci. Disaggregated of results by B) Conservative temperature (ºC) C) Oxygen concentration (mL/L), D) Nitrate concentration (µM), E) Absolute salinity (g/Kg), F) Phosphate concentration (µM) and G) Silicate (µM) concentration. The set size (horizontal bars) indicates the total number of loci correlated with that environmental variable. The intersection size (vertical bars) indicates the number of loci per set intersection, the dots represent the set of unique loci correlated only with that environmental variable and no other one, while the dots connected by line represent loci shared by two, three, or more environmental variable.

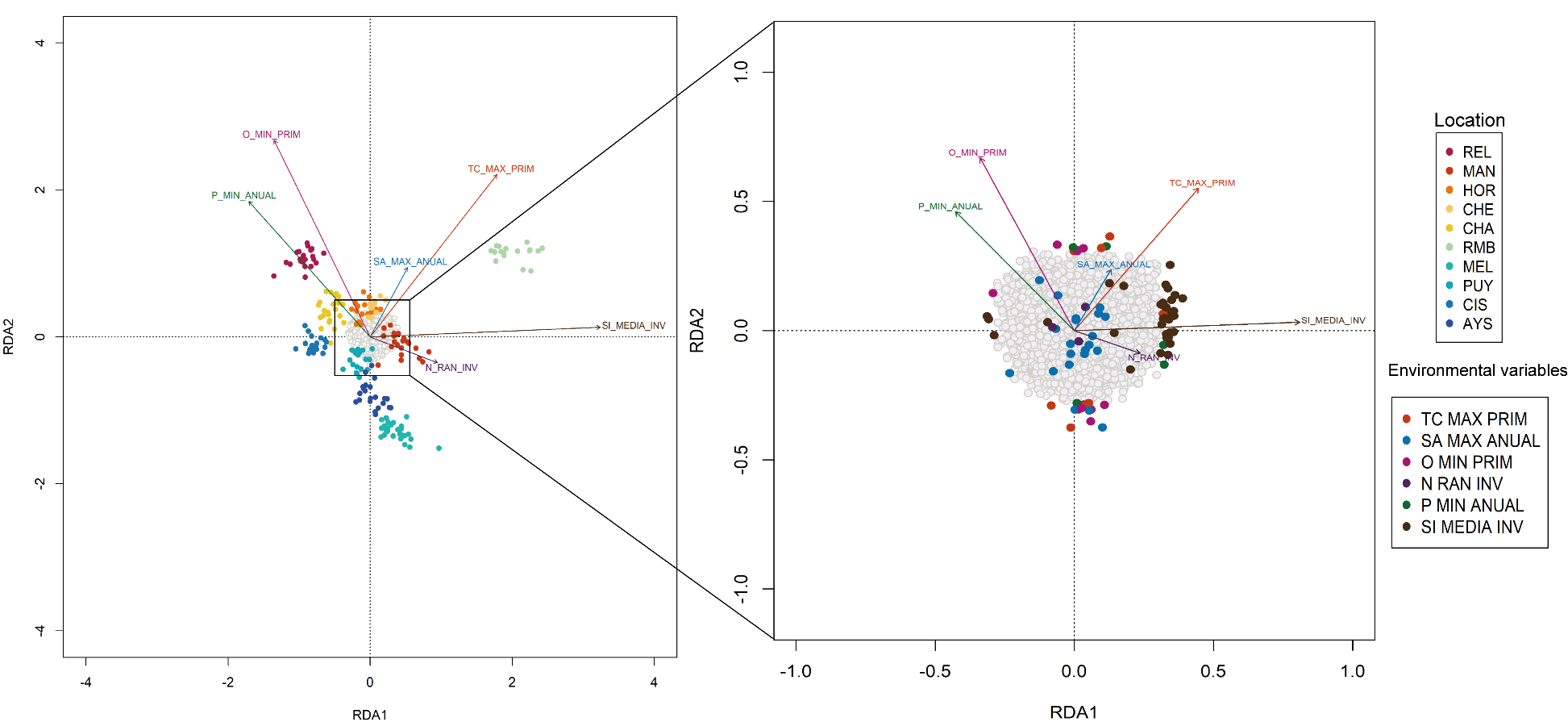

Figure S 7 Redundancy analysis (*RDA*) using the 11,961 loci.

Arrows represent environmental variables (TC_MAX_PRIM: Spring maximum temperature, SA_MAX_ANUAL: Annual maximum salinity, O_MIN_PRIM: Spring minimum oxygen, P_MIN_ANUAL: Annual minimum phosphate, N_RAN_INV: winter nitrate range and SI_MEDIA_INV: Winter average silicate). A) Plots show the distribution of loci (gray dots) and genotypes of individuals (colored circles) are plotted according to their sampling location. B) Putative adaptative loci (colored circles) are plotted according to their predictor (environmental variables).

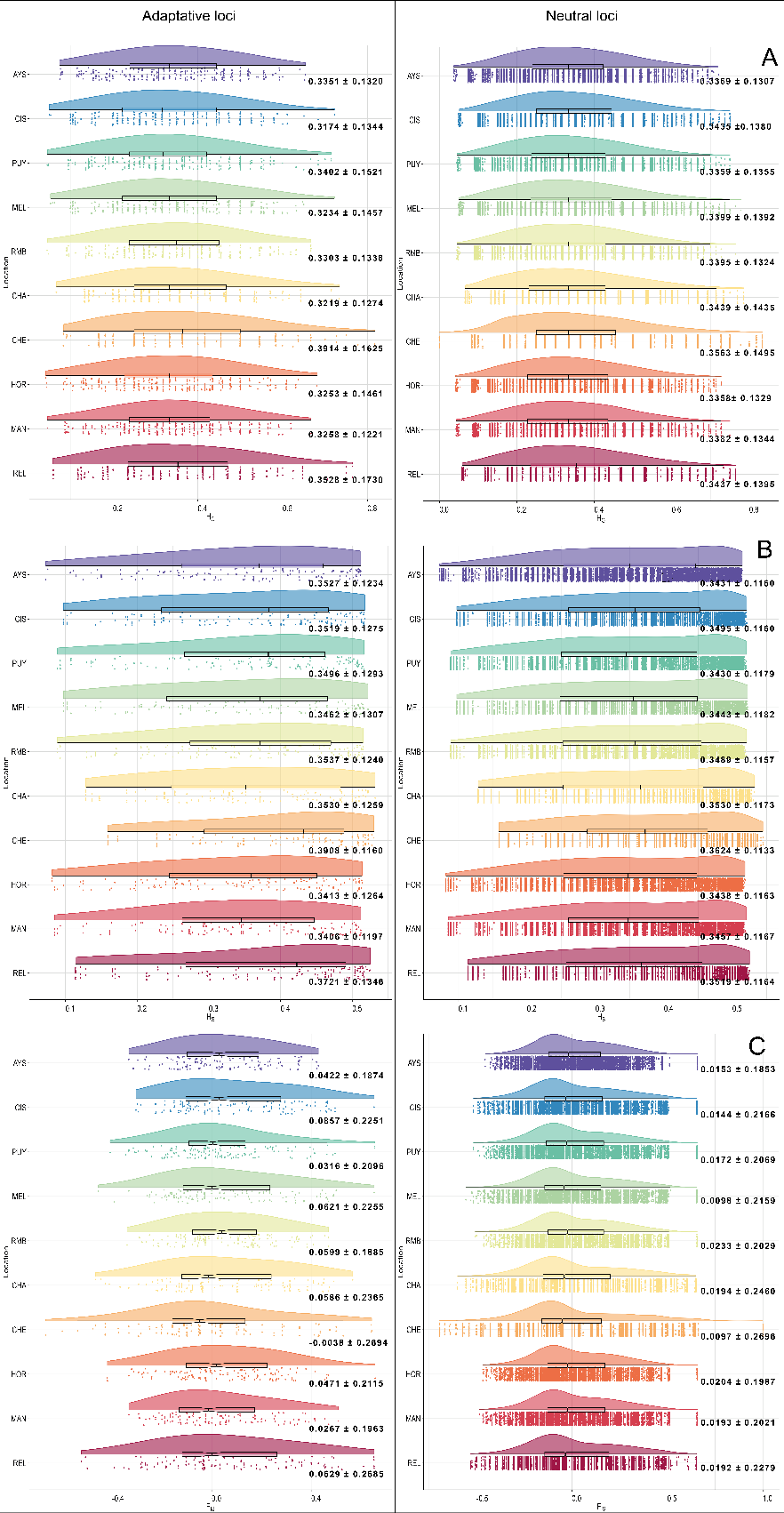

Figure S 8 Raincloud plots: Distribution of estimates of genetic diversity.

A) H_O_ = observed heterozygosity, B) H_E_ = expected heterozygosity and C) F_IS_ inbreeding coefficient. Index was calculated using 131 loci for adaptive dataset and 9,536 loci for neutral dataset.

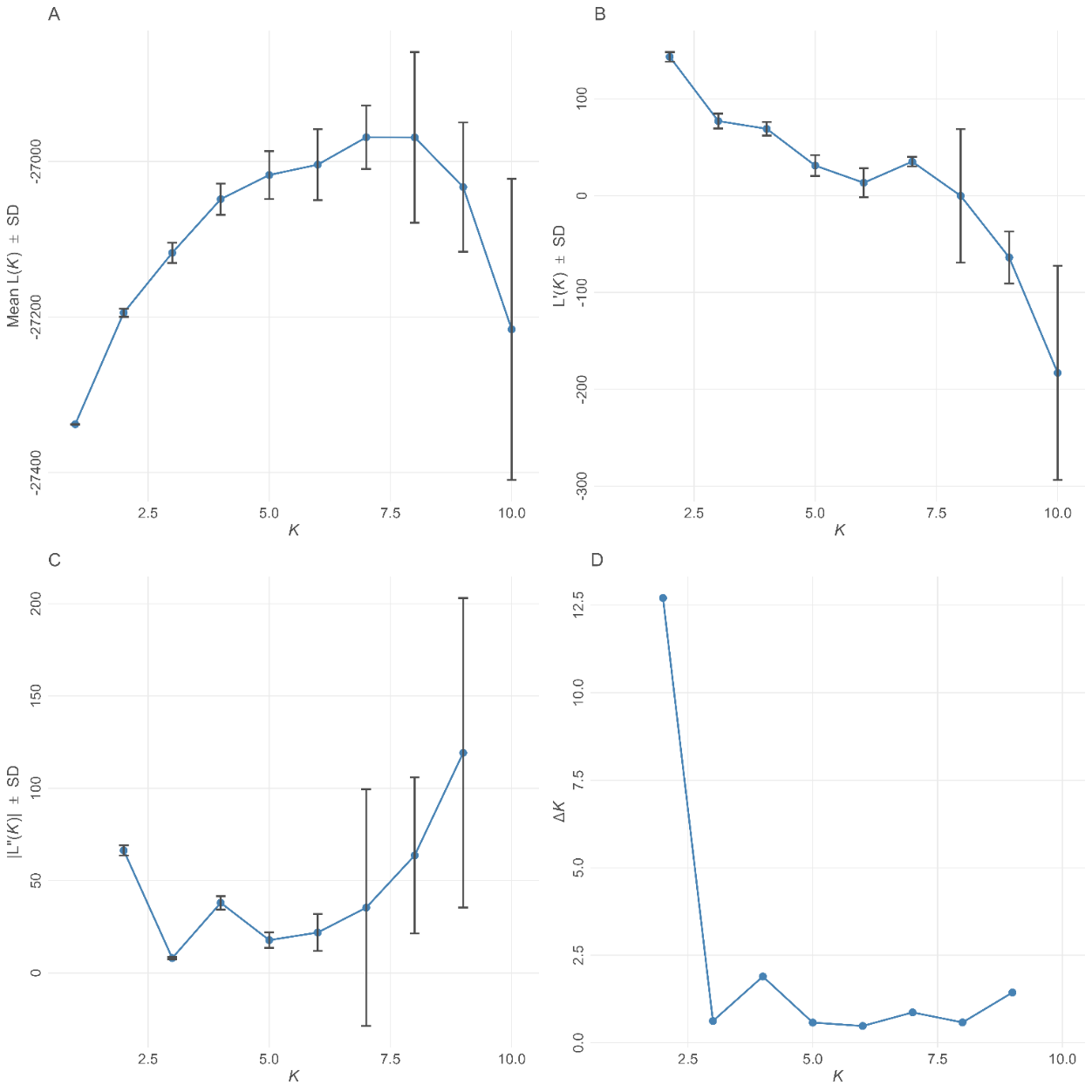

Figure S 9 Estimates of the number of clusters (K) for adaptive loci.

Those using different statistics using the Evanno method to determine the ideal number of clusters by putatively adaptative loci. Error bars are standard deviation. A) Estimated log probability of the data for the 10 Structure runs at each K, B) first derivative, C) second derivative and D) ΔK, the rate of change in the log probability of data between successive K values.

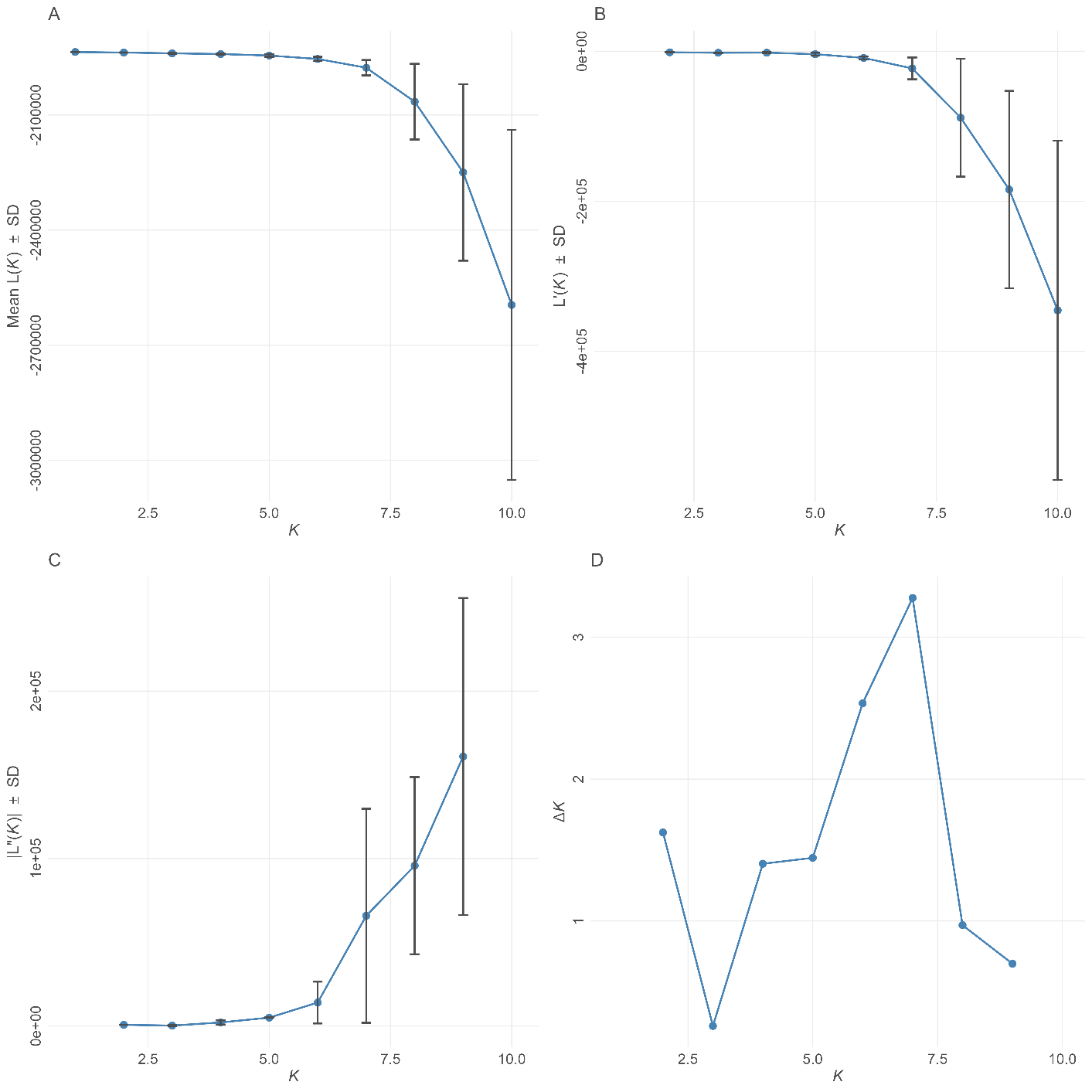

Figure S 10 Estimates of the number of clusters (K) for neutral loci.

using different statistics using the Evanno method to determine the ideal number of clusters by putatively neutral loci. Error bars are standard deviation. A) Estimated log probability of the data for the 10 Structure runs at each K, B) first derivative, C) second derivative and D) ΔK, the rate of change in the log probability of data between successive K values.

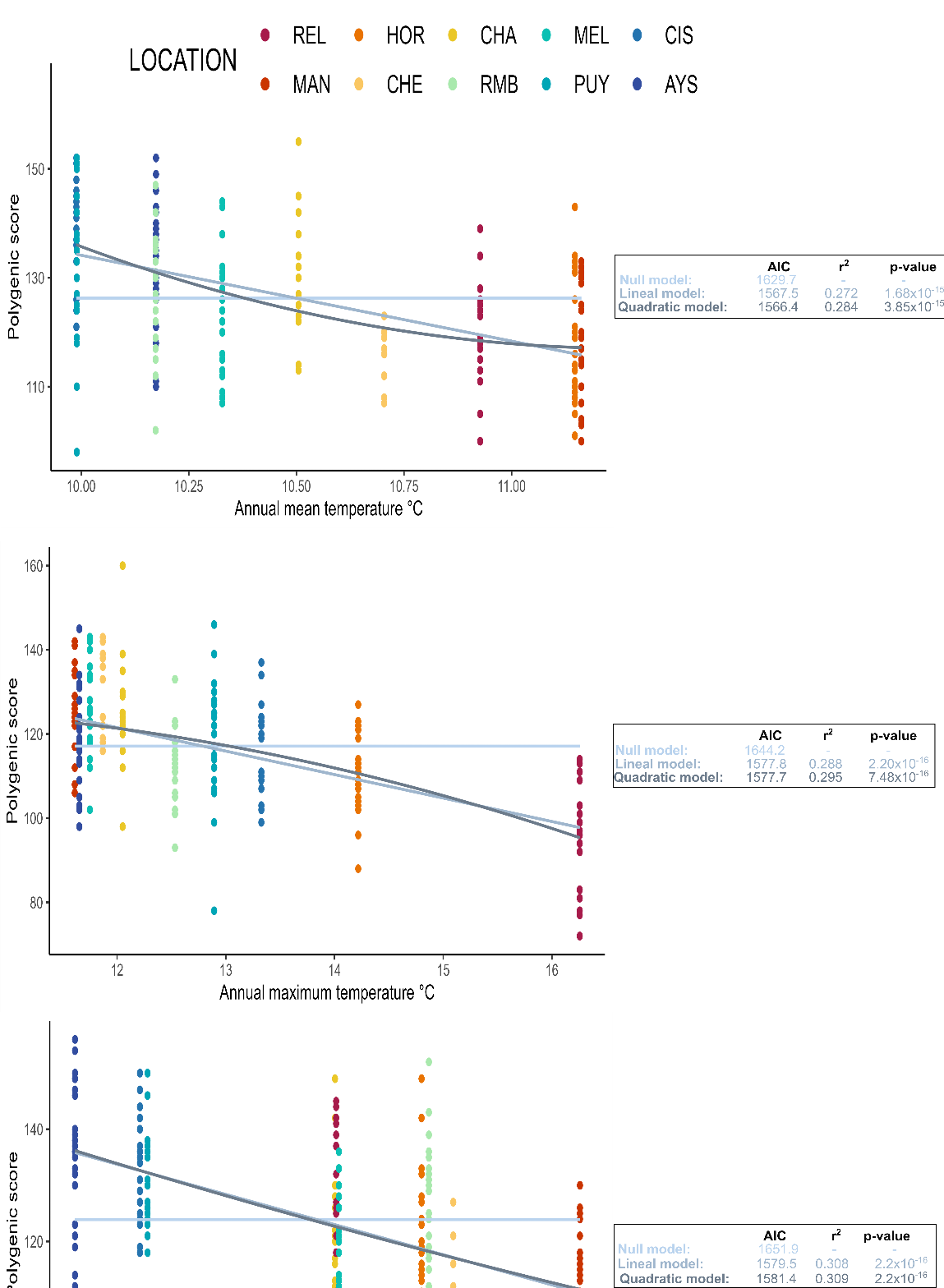

Figure S 11 Correlations between additive polygenic scores (APS) based on A) Annual mean temperature B) Annual maximum temperature and C) Annual minimum temperature and 131 putative adaptative loci. Correlation coefficient (R^2^) and p-values and AIC are presented for each variable.

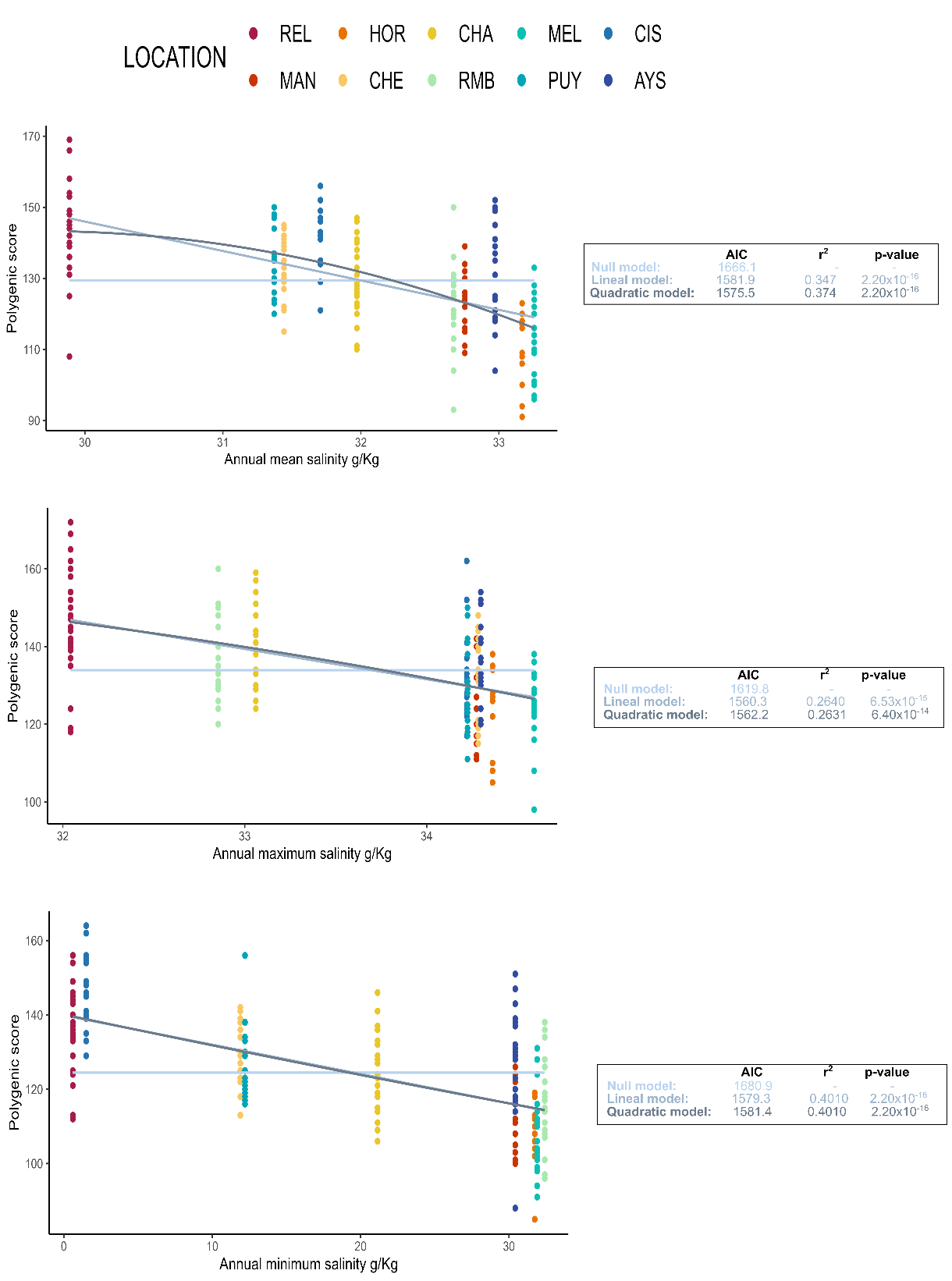

Figure S 12 Correlations between additive polygenic scores (APS) based on A) Annual mean salinity B) Annual maximum salinity and C) Annual minimum salinity and 131 putative adaptative loci. Correlation coefficient (R^2^) and p-values and AIC are presented for each variable.

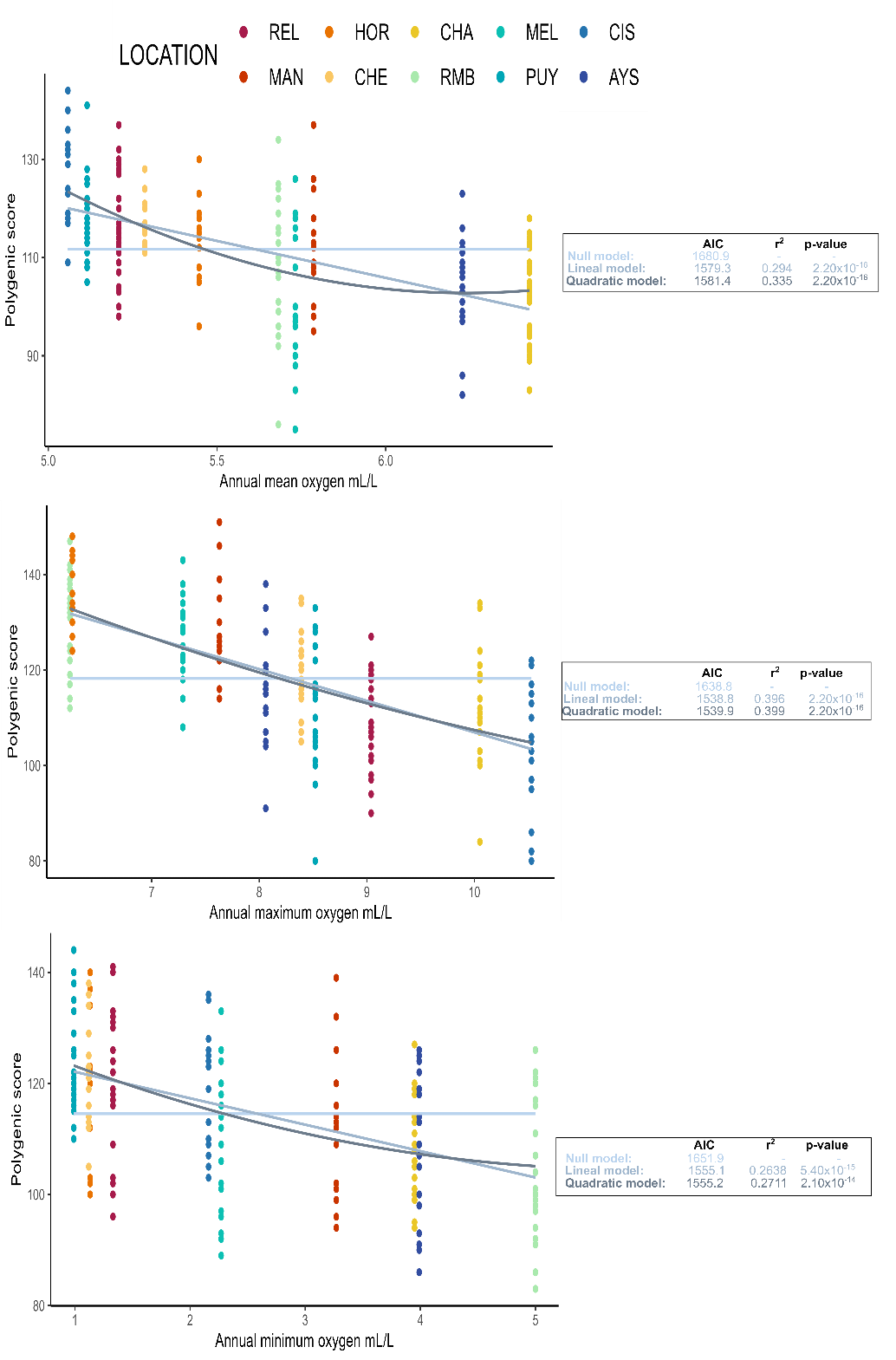

Figure S 13 Correlations between additive polygenic scores (APS) based on A) Annual mean oxygen concentration B) Annual maximum oxygen concentration and C) Annual minimum oxygen concentration and 131 putative adaptative loci. Correlation coefficient (R2) and p-values and AIC are presented for each variable.

#### Supplementary Tables

Table S 1 Details of environmental variables for landscape genomics analysis.

| **Location ID** | **Latitude** | **Longitude** | **Annual conservative temperature (ºC)** | | | | **Annual absolute salinity (g Kg^-1^)** | | | | **Annual oxygen concentration (mL L^-1^)** | | | |
| --- | --- | --- | --- | --- | --- | --- | --- | --- | --- | --- | --- | --- | --- | --- |
|  |  |  | **Mean** | **Min.** | **Max.** | **Range** | **Mean** | **Min.** | **Max.** | **Range** | **Mean** | **Min.** | **Max.** | **Range** |
| REL | -41.533111 | -72.302351 | 10.927 | 8.278 | 16.256 | 7.978 | 31.707 | 1.508 | 34.220 | 32.712 | 5.058 | 2.160 | 10.530 | 8.370 |
| MAN | -41.880830 | -73.516390 | 11.162 | 10.738 | 11.611 | 0.873 | 32.671 | 32.411 | 32.854 | 0.444 | 5.682 | 5.000 | 6.240 | 1.240 |
| HOR | -41.969422 | -72.444251 | 11.147 | 9.140 | 14.218 | 5.078 | 31.971 | 21.143 | 33.061 | 11.918 | 6.426 | 3.950 | 10.050 | 6.100 |
| CHE * | -42.042500 | -74.032778 | 10.704 | 9.460 | 11.869 | 2.409 | 33.168 | 31.735 | 34.363 | 2.627 | 5.447 | 1.132 | 6.263 | 5.131 |
| CHA | -42.897317 | -72.737594 | 10.505 | 8.267 | 12.051 | 3.784 | 32.753 | 30.422 | 34.274 | 3.852 | 5.786 | 3.270 | 7.630 | 4.360 |
| RMB | -43.744083 | -72.978328 | 10.173 | 9.214 | 12.533 | 3.319 | 32.972 | 30.420 | 34.298 | 3.877 | 6.227 | 3.990 | 8.060 | 4.070 |
| MEL | -43.876376 | -73.891024 | 10.328 | 8.306 | 11.750 | 3.444 | 33.256 | 31.901 | 34.591 | 2.690 | 5.732 | 2.270 | 7.290 | 5.020 |
| PUY | -44.326764 | -72.562060 | 9.990 | 6.376 | 12.893 | 6.517 | 31.372 | 12.210 | 34.224 | 22.014 | 5.115 | 0.990 | 8.520 | 7.530 |
| CIS | -44.743602 | -72.699889 | 9.989 | 6.299 | 13.327 | 7.028 | 31.443 | 11.896 | 34.284 | 22.388 | 5.286 | 1.120 | 8.390 | 7.270 |
| AYS | -45.250000 | -73.250000 | 10.174 | 5.644 | 11.653 | 6.009 | 29.888 | 0.608 | 32.043 | 31.435 | 5.209 | 1.330 | 9.040 | 7.710 |

| **Location ID** | **Latitude** | **Longitude** | **Annual nitrate concentration (µM)** | | | | **Annual phosphate concentration (µM)** | | | | **Annual silicate concentration (µM)** | | | |
| --- | --- | --- | --- | --- | --- | --- | --- | --- | --- | --- | --- | --- | --- | --- |
|  |  |  | **Mean** | **Min.** | **Max.** | **Range** | **Mean** | **Min.** | **Max.** | **Range** | **Mean** | **Min.** | **Max.** | **Range** |
| REL | -41.533111 | -72.302351 | 18.681 | 0.000 | 35.000 | 35.000 | 1.846 | 0.000 | 3.150 | 3.150 | 35.688 | 0.000 | 210.779 | 210.779 |
| MAN | -41.880830 | -73.516390 | 16.440 | 6.910 | 23.060 | 16.150 | 1.748 | 0.870 | 2.220 | 1.350 | 12.772 | 6.320 | 53.320 | 47.000 |
| HOR | -41.969422 | -72.444251 | 11.351 | 0.000 | 24.700 | 24.700 | 1.391 | 0.050 | 2.720 | 2.670 | 10.614 | 0.000 | 29.000 | 29.000 |
| CHE * | -42.042500 | -74.032778 | 11.187 | 0.600 | 32.800 | 32.200 | 1.102 | 0.230 | 2.750 | 2.520 | 5.750 | 0.000 | 21.000 | 21.000 |
| CHA | -42.897317 | -72.737594 | 17.534 | 3.300 | 27.200 | 23.900 | 1.758 | 0.840 | 2.780 | 1.940 | 12.735 | 0.330 | 76.000 | 75.670 |
| RMB | -43.744083 | -72.978328 | 11.718 | 0.000 | 18.460 | 18.460 | 1.291 | 0.620 | 1.990 | 1.370 | 7.836 | 1.190 | 15.570 | 14.380 |
| MEL | -43.876376 | -73.891024 | 13.769 | 0.074 | 29.100 | 29.026 | 1.384 | 0.540 | 2.600 | 2.060 | 8.409 | 0.000 | 19.000 | 19.000 |
| PUY | -44.326764 | -72.562060 | 16.271 | 0.000 | 29.257 | 29.257 | 1.493 | 0.000 | 3.620 | 3.620 | 16.163 | 1.000 | 77.000 | 76.000 |
| CIS | -44.743602 | -72.699889 | 15.079 | 0.000 | 30.945 | 30.945 | 1.407 | 0.000 | 3.530 | 3.530 | 16.913 | 1.000 | 63.000 | 62.000 |
| AYS | -45.250000 | -73.250000 | 16.017 | 0.000 | 28.400 | 28.400 | 1.416 | 0.000 | 2.570 | 2.570 | 30.479 | 3.000 | 181.000 | 178.000 |

| **Location ID** | **Latitude** | **Longitude** | **Spring conservative temperature (ºC)** | | | | **Spring absolute salinity (g Kg^-1^)** | | | | **Spring oxygen concentration (mL L^-1^)** | | | |
| --- | --- | --- | --- | --- | --- | --- | --- | --- | --- | --- | --- | --- | --- | --- |
|  |  |  | **Mean** | **Min.** | **Max.** | **Range** | **Mean** | **Min.** | **Max.** | **Range** | **Mean** | **Min.** | **Max.** | **Range** |
| REL | -41.533111 | -72.302351 | 10.716 | 8.331 | 16.256 | 7.925 | 31.306 | 1.508 | 34.220 | 32.712 | 5.736 | 2.160 | 10.350 | 8.190 |
| MAN | -41.880830 | -73.516390 | 11.263 | 10.897 | 13.351 | 2.454 | 32.660 | 32.020 | 32.789 | 0.769 | 5.984 | 5.160 | 8.030 | 2.870 |
| HOR | -41.969422 | -72.444251 | 11.200 | 10.012 | 14.218 | 4.206 | 31.941 | 21.143 | 32.996 | 11.852 | 6.428 | 4.110 | 10.050 | 5.940 |
| CHE * | -42.042500 | -74.032778 | 10.776 | 10.499 | 11.056 | 0.557 | 33.331 | 33.038 | 33.864 | 0.826 | 5.850 | 5.042 | 6.191 | 1.149 |
| CHA | -42.897317 | -72.737594 | 10.524 | 8.267 | 12.051 | 3.784 | 32.817 | 31.494 | 34.274 | 2.780 | 5.769 | 3.270 | 7.630 | 4.360 |
| RMB | -43.744083 | -72.978328 | 10.310 | 9.214 | 12.533 | 3.319 | 32.647 | 30.420 | 34.298 | 3.877 | 6.321 | 3.990 | 8.060 | 4.070 |
| MEL | -43.876376 | -73.891024 | 10.149 | 8.306 | 11.314 | 3.008 | 33.299 | 31.413 | 34.591 | 3.178 | 5.672 | 2.310 | 7.290 | 4.980 |
| PUY | -44.326764 | -72.562060 | 10.113 | 8.881 | 12.893 | 4.012 | 31.659 | 12.210 | 34.224 | 22.014 | 5.283 | 0.990 | 8.520 | 7.530 |
| CIS | -44.743602 | -72.699889 | 10.122 | 8.806 | 13.327 | 4.521 | 31.721 | 11.896 | 34.284 | 22.388 | 5.595 | 1.120 | 8.130 | 7.010 |
| AYS | -45.250000 | -73.250000 | 10.168 | 8.687 | 11.653 | 2.966 | 29.602 | 0.608 | 32.151 | 31.544 | 5.485 | 1.580 | 8.510 | 6.930 |

|  | **Location ID** | | **Latitude** | | **Longitude** | | **Spring nitrate concentration (µM)** | | | | | | | | | **Spring phosphate concentration (µM)** | | | | | | | **Spring silicate concentration (µM)** | | | | | | | |
| --- | --- | --- | --- | --- | --- | --- | --- | --- | --- | --- | --- | --- | --- | --- | --- | --- | --- | --- | --- | --- | --- | --- | --- | --- | --- | --- | --- | --- | --- | --- |
|  |  |  |  |  |  |  | **Mean** | | **Min.** | **Max.** | | | **Range** | | | **Mean** | | **Min.** | **Max.** | | **Range** | | **Mean** | | **Min.** | | **Max.** | | | **Range** |
|  | REL | | -41.533111 | | -72.302351 | | 15.373 | | 0.000 | 29.200 | | | 29.200 | | | 1.555 | | 0.000 | 3.150 | | 3.150 | | 25.855 | | 0.000 | | 112.000 | | | 112.000 |
|  | MAN | | -41.880830 | | -73.516390 | | 16.176 | | 0.410 | 20.160 | | | 19.750 | | | 1.782 | | 0.560 | 2.220 | | 1.660 | | 13.129 | | 1.360 | | 53.320 | | | 51.960 |
|  | HOR | | -41.969422 | | -72.444251 | | 8.712 | | 0.000 | 24.700 | | | 24.700 | | | 1.134 | | 0.050 | 2.720 | | 2.670 | | 7.710 | | 0.000 | | 28.860 | | | 28.860 |
|  | CHE * | | -42.042500 | | -74.032778 | | 16.926 | | 0.600 | 32.800 | | | 32.200 | | | 1.362 | | 0.230 | 2.630 | | 2.400 | | 7.905 | | 0.000 | | 19.000 | | | 19.000 |
|  | CHA | | -42.897317 | | -72.737594 | | 15.946 | | 3.300 | 27.200 | | | 23.900 | | | 1.633 | | 0.840 | 2.780 | | 1.940 | | 10.077 | | 0.330 | | 45.000 | | | 44.670 |
|  | RMB | | -43.744083 | | -72.978328 | | 9.917 | | 0.000 | 18.460 | | | 18.460 | | | 1.237 | | 0.620 | 1.990 | | 1.370 | | 5.963 | | 1.190 | | 12.000 | | | 10.810 |
|  | MEL | | -43.876376 | | -73.891024 | | 12.003 | | 0.074 | 28.700 | | | 28.626 | | | 1.325 | | 0.640 | 2.310 | | 1.670 | | 6.361 | | 0.000 | | 17.000 | | | 17.000 |
|  | PUY | | -44.326764 | | -72.562060 | | 13.203 | | 0.000 | 28.900 | | | 28.900 | | | 1.283 | | 0.000 | 3.620 | | 3.620 | | 17.080 | | 1.000 | | 77.000 | | | 76.000 |
|  | CIS | | -44.743602 | | -72.699889 | | 10.343 | | 0.000 | 29.100 | | | 29.100 | | | 1.058 | | 0.000 | 3.530 | | 3.530 | | 17.104 | | 1.000 | | 63.000 | | | 62.000 |
|  | AYS | | -45.250000 | | -73.250000 | | 13.248 | | 0.000 | 25.100 | | | 25.100 | | | 1.253 | | 0.000 | 2.570 | | 2.570 | | 28.964 | | 3.000 | | 105.000 | | | 102.000 |
| **Location ID** | | **Latitude** | | **Longitude** | | **Winter conservative temperature (ºC)** | | | | | | | | | **Winter absolute salinity (g Kg^-1^)** | | | | | | | | | **Winter oxygen concentration (mL L^-1^)** | | | | | | |
|  |  |  |  |  |  | **Mean** | | **Min.** | | | **Max.** | **Range** | | **Mean** | | | **Min.** | | | **Max.** | | **Range** | | **Mean** | | **Min.** | | **Max.** | **Range** | |
| REL  MAN | | -41.533111  -41.880830 | | -72.302351  -73.516390 | | 11.168  10.842 | | 8.278  10.738 | | | 11.626  10.974 | 3.348  0.236 | | 31.827  32.644 | | | 2.546  32.411 | | | 33.221  32.854 | | 30.675  0.444 | | 4.595  5.697 | | 2.210  5.000 | | 10.530  6.170 | 8.320  1.170 | |
| HOR | | -41.969422 | | -72.444251 | | 10.815 | | 9.140 | | | 11.493 | 2.353 | | 32.256 | | | 27.932 | | | 33.061 | | 5.129 | | 6.488 | | 3.950 | | 9.440 | 5.490 | |
| CHE * | | -42.042500 | | -74.032778 | | 10.696 | | 9.460 | | | 11.869 | 2.409 | | 33.152 | | | 31.735 | | | 34.452 | | 2.717 | | 5.407 | | 1.132 | | 6.263 | 5.131 | |
| CHA | | -42.897317 | | -72.737594 | | 10.318 | | 9.664 | | | 10.571 | 0.907 | | 32.579 | | | 30.422 | | | 33.303 | | 2.881 | | 5.985 | | 5.140 | | 7.550 | 2.410 | |
| RMB | | -43.744083 | | -72.978328 | | 10.294 | | 10.242 | | | 10.394 | 0.152 | | 32.853 | | | 32.746 | | | 33.069 | | 0.323 | | 6.000 | | 5.550 | | 6.190 | 0.640 | |
| MEL | | -43.876376 | | -73.891024 | | 10.472 | | 9.138 | | | 11.750 | 2.612 | | 33.180 | | | 32.168 | | | 34.402 | | 2.234 | | 5.904 | | 2.270 | | 6.650 | 4.380 | |
| PUY | | -44.326764 | | -72.562060 | | 9.816 | | 6.376 | | | 10.606 | 4.230 | | 31.111 | | | 14.577 | | | 34.197 | | 19.620 | | 5.010 | | 1.540 | | 8.088 | 6.548 | |
| CIS | | -44.743602 | | -72.699889 | | 9.662 | | 6.299 | | | 10.606 | 4.307 | | 31.312 | | | 13.800 | | | 34.226 | | 20.426 | | 5.338 | | 1.540 | | 8.390 | 6.850 | |
| AYS | | -45.250000 | | -73.250000 | | 10.188 | | 5.644 | | | 11.249 | 5.605 | | 30.131 | | | 4.483 | | | 32.043 | | 27.560 | | 5.059 | | 1.330 | | 9.040 | 7.710 | |

| **Location ID** | **Latitude** | **Longitude** | **Winter nitrate concentration (µM)** | | | | **Winter phosphate concentration (µM)** | | | | **Winter silicate concentration (µM)** | | | |
| --- | --- | --- | --- | --- | --- | --- | --- | --- | --- | --- | --- | --- | --- | --- |
|  |  |  | **Mean** | **Min.** | **Max.** | **Range** | **Mean** | **Min.** | **Max.** | **Range** | **Mean** | **Min.** | **Max.** | **Range** |
| REL | -41.533111 | -72.302351 | 21.339 | 0.590 | 35.000 | 34.410 | 2.084 | 0.000 | 3.038 | 3.038 | 44.173 | 1.330 | 210.779 | 209.449 |
| MAN | -41.880830 | -73.516390 | 18.480 | 6.910 | 23.060 | 16.150 | 1.553 | 0.870 | 1.850 | 0.980 | 16.440 | 10.410 | 22.190 | 11.780 |
| HOR | -41.969422 | -72.444251 | 15.696 | 0.000 | 22.900 | 22.900 | 1.800 | 0.190 | 2.590 | 2.400 | 15.333 | 0.000 | 29.000 | 29.000 |
| CHE * | -42.042500 | -74.032778 | 23.128 | 11.200 | 32.500 | 21.300 | 2.061 | 1.160 | 2.750 | 1.590 | 13.240 | 3.000 | 21.000 | 18.000 |
| CHA | -42.897317 | -72.737594 | 19.979 | 7.700 | 23.774 | 16.074 | 1.951 | 0.910 | 2.280 | 1.370 | 18.363 | 3.830 | 76.000 | 72.170 |
| RMB | -43.744083 | -72.978328 | 17.688 | 17.570 | 17.750 | 0.180 | 1.390 | 1.310 | 1.440 | 0.130 | 13.475 | 12.470 | 15.570 | 3.100 |
| MEL | -43.876376 | -73.891024 | 15.204 | 5.240 | 29.100 | 23.860 | 1.416 | 0.540 | 2.600 | 2.060 | 10.166 | 2.880 | 19.000 | 16.120 |
| PUY | -44.326764 | -72.562060 | 19.572 | 5.139 | 29.257 | 24.118 | 1.692 | 0.127 | 3.100 | 2.973 | 16.498 | 2.690 | 43.431 | 40.741 |
| CIS | -44.743602 | -72.699889 | 19.093 | 0.900 | 30.945 | 30.045 | 1.659 | 0.085 | 3.100 | 3.015 | 17.216 | 4.353 | 43.431 | 39.078 |
| AYS | -45.250000 | -73.250000 | 18.664 | 2.100 | 28.400 | 26.300 | 1.564 | 0.000 | 2.550 | 2.550 | 30.596 | 9.000 | 181.000 | 172.000 |

Note 1. Summary statistics for environmental variables are based on data collected from CIMAR-FIORDOS 1995-2018 at depths between 0 and 100 m.

Note 2. The information of CHE* was supplemented with data provided by IFOP (https://www.ifop.cl/chonos/) and Bio-ORACLE (https://www.bio-oracle.org)

Table S 2 Linear regression statistics for each of the adaptative loci.

Those that showing strong correlation (r > 0.60) with an environmental variable

| **Loci ID** | **R^2^** | **F** | **p-value** | **p-value _adjusted_** | **Environmental variable** |
| --- | --- | --- | --- | --- | --- |
| 5767_257 | 0.67 | 16.44 | 0.0037 | 0.0089 | Annual mean temperature |
| 5767_257 | 0.67 | 15.90 | 0.0040 | 0.0089 | Spring mean temperature |
| 48326_234 | 0.76 | 24.77 | 0.0011 | 0.0074 | Winter mean temperature |
| 54816_33 | 0.66 | 15.26 | 0.0045 | 0.0089 | Winter mean temperature |
| 18407_279 | 0.71 | 19.35 | 0.0023 | 0.0089 | Annual maximum temperature |
| 8969_106 | 0.65 | 15.00 | 0.0047 | 0.0089 | Annual maximum temperature |
| 58479_12 | 0.65 | 14.53 | 0.0051 | 0.0089 | Annual maximum temperature |
| 41766_287 | 0.63 | 13.64 | 0.0061 | 0.0089 | Annual maximum temperature |
| 37756_46 | 0.66 | 15.67 | 0.0042 | 0.0089 | Winter maximum temperature |
| 50745_175 | 0.64 | 14.31 | 0.0054 | 0.0089 | Winter maximum temperature |
| 39206_170 | 0.68 | 17.18 | 0.0032 | 0.0089 | Annual minimum temperature |
| 43097_144 | 0.67 | 15.96 | 0.0040 | 0.0089 | Annual minimum temperature |
| 18596_155 | 0.66 | 15.63 | 0.0042 | 0.0089 | Annual minimum temperature |
| 55344_89 | 0.60 | 11.86 | 0.0088 | 0.0095 | Annual minimum temperature |
| 39206_170 | 0.74 | 23.15 | 0.0013 | 0.0074 | Winter minimum temperature |
| 50669_80 | 0.61 | 12.33 | 0.0079 | 0.0092 | Winter minimum temperature |
| 18407_279 | 0.83 | 38.43 | 0.0003 | 0.0056 | Annual temperature range |
| 36199_80 | 0.62 | 13.26 | 0.0066 | 0.0089 | Annual temperature range |
| 4598_98 | 0.77 | 27.43 | 0.0008 | 0.0074 | Spring temperature range |
| 18407_279 | 0.66 | 15.60 | 0.0042 | 0.0089 | Spring temperature range |
| 52788_248 | 0.65 | 14.76 | 0.0049 | 0.0089 | Spring temperature range |
| 15060_124 | 0.62 | 13.25 | 0.0066 | 0.0089 | Spring temperature range |
| 39206_170 | 0.78 | 28.17 | 0.0007 | 0.0074 | Winter temperature range |
| 36199_80 | 0.60 | 12.19 | 0.0082 | 0.0092 | Annual minimum salinity |
| 36199_80 | 0.62 | 13.08 | 0.0068 | 0.0089 | Spring minimum salinity |
| 18407_279 | 0.60 | 11.68 | 0.0091 | 0.0095 | Winter minimum salinity |
| 16449_9 | 0.74 | 22.37 | 0.0015 | 0.0074 | Annual maximum salinity |
| 16449_9 | 0.67 | 16.28 | 0.0038 | 0.0089 | Spring maximum salinity |
| 6629_190 | 0.61 | 12.73 | 0.0073 | 0.0089 | Spring maximum salinity |
| 16449_9 | 0.61 | 12.45 | 0.0078 | 0.0091 | Winter maximum salinity |
| 18407_279 | 0.63 | 13.41 | 0.0064 | 0.0089 | Annual salinity range |
| 36199_80 | 0.60 | 11.61 | 0.0093 | 0.0095 | Annual salinity range |
| 36199_80 | 0.61 | 12.74 | 0.0073 | 0.0089 | Spring salinity range |
| 18407_279 | 0.61 | 12.63 | 0.0075 | 0.0089 | Spring salinity range |
| 18407_279 | 0.63 | 13.62 | 0.0061 | 0.0089 | Winter salinity range |
| 50669_80 | 0.75 | 24.11 | 0.0012 | 0.0074 | Spring mean oxygen concentration |
| 39206_170 | 0.63 | 13.35 | 0.0065 | 0.0089 | Spring mean oxygen concentration |
| 22359_124 | 0.84 | 42.42 | 0.0002 | 0.0056 | Annual maximum oxygen concentration |
| 37533_59 | 0.66 | 15.51 | 0.0043 | 0.0089 | Annual maximum oxygen concentration |
| 36199_80 | 0.61 | 12.31 | 0.0080 | 0.0092 | Annual maximum oxygen concentration |
| 58479_12 | 0.60 | 11.59 | 0.0093 | 0.0095 | Annual maximum oxygen concentration |
| 22359_124 | 0.68 | 17.27 | 0.0032 | 0.0089 | Spring maximum oxygen concentration |
| 58479_12 | 0.67 | 16.16 | 0.0038 | 0.0089 | Spring maximum oxygen concentration |
| 37533_59 | 0.64 | 14.45 | 0.0052 | 0.0089 | Spring maximum oxygen concentration |
| 15060_124 | 0.63 | 13.65 | 0.0061 | 0.0089 | Spring maximum oxygen concentration |
| 13680_161 | 0.61 | 12.75 | 0.0073 | 0.0089 | Spring maximum oxygen concentration |
| 36199_80 | 0.60 | 12.18 | 0.0082 | 0.0092 | Spring maximum oxygen concentration |
| 36199_80 | 0.75 | 24.31 | 0.0011 | 0.0074 | Winter maximum oxygen concentration |
| 22359_124 | 0.73 | 21.42 | 0.0017 | 0.0074 | Winter maximum oxygen concentration |
| 18407_279 | 0.69 | 18.01 | 0.0028 | 0.0089 | Winter maximum oxygen concentration |
| 50000_74 | 0.69 | 17.42 | 0.0031 | 0.0089 | Winter maximum oxygen concentration |
| 58479_12 | 0.63 | 13.41 | 0.0064 | 0.0089 | Winter maximum oxygen concentration |
| 39206_170 | 0.60 | 11.51 | 0.0095 | 0.0095 | Spring minimum oxygen concentration |
| 63849_179 | 0.62 | 13.31 | 0.0065 | 0.0089 | Winter minimum oxygen concentration |
| 39206_170 | 0.60 | 11.48 | 0.0095 | 0.0095 | Winter minimum oxygen concentration |
| 18407_279 | 0.75 | 23.87 | 0.0012 | 0.0074 | Annual oxygen concentration range |
| 36199_80 | 0.77 | 26.99 | 0.0008 | 0.0074 | Spring oxygen concentration range |
| 18407_279 | 0.62 | 12.92 | 0.0070 | 0.0089 | Spring oxygen concentration range |
| 18407_279 | 0.66 | 15.71 | 0.0042 | 0.0089 | Winter oxygen concentration range |
| 54816_33 | 0.60 | 11.71 | 0.0091 | 0.0095 | Spring mean nitrate concentration |
| 9580_78 | 0.63 | 13.41 | 0.0064 | 0.0089 | Winter mean nitrate concentration |
| 63849_179 | 0.60 | 12.18 | 0.0082 | 0.0092 | Spring maximum nitrate concentration |
| 43097_144 | 0.65 | 14.85 | 0.0049 | 0.0089 | Annual minimum nitrate concentration |
| 19396_225 | 0.80 | 32.31 | 0.0005 | 0.0074 | Spring minimum nitrate concentration |
| 63106_152 | 0.74 | 22.75 | 0.0014 | 0.0074 | Spring minimum nitrate concentration |
| 20305_146 | 0.73 | 21.75 | 0.0016 | 0.0074 | Spring minimum nitrate concentration |
| 53488_180 | 0.73 | 21.20 | 0.0017 | 0.0074 | Spring minimum nitrate concentration |
| 27224_149 | 0.73 | 21.10 | 0.0018 | 0.0074 | Spring minimum nitrate concentration |
| 51829_60 | 0.67 | 16.05 | 0.0039 | 0.0089 | Spring minimum nitrate concentration |
| 1583_131 | 0.64 | 13.92 | 0.0058 | 0.0089 | Spring minimum nitrate concentration |
| 13102_202 | 0.63 | 13.69 | 0.0060 | 0.0089 | Spring minimum nitrate concentration |
| 36199_80 | 0.83 | 38.11 | 0.0003 | 0.0056 | Winter minimum nitrate concentration |
| 50000_74 | 0.65 | 14.96 | 0.0048 | 0.0089 | Winter minimum nitrate concentration |
| 18407_279 | 0.62 | 12.81 | 0.0072 | 0.0089 | Annual nitrate concentration range |
| 63849_179 | 0.62 | 13.27 | 0.0066 | 0.0089 | Spring nitrate concentration range |
| 37301_273 | 0.63 | 13.57 | 0.0062 | 0.0089 | Annual mean phosphate concentration |
| 43097_144 | 0.63 | 13.43 | 0.0064 | 0.0089 | Spring mean phosphate concentration |
| 54816_33 | 0.60 | 11.80 | 0.0089 | 0.0095 | Spring mean phosphate concentration |
| 18407_279 | 0.60 | 11.60 | 0.0093 | 0.0095 | Spring maximum phosphate concentration |
| 18407_279 | 0.61 | 12.66 | 0.0074 | 0.0089 | Winter maximum phosphate concentration |
| 50000_74 | 0.63 | 13.40 | 0.0064 | 0.0089 | Annual minimum phosphate concentration |
| 50000_74 | 0.73 | 21.22 | 0.0017 | 0.0074 | Spring minimum phosphate concentration |
| 78679_308 | 0.64 | 14.11 | 0.0056 | 0.0089 | Spring minimum phosphate concentration |
| 60830_31 | 0.60 | 12.08 | 0.0084 | 0.0092 | Spring minimum phosphate concentration |
| 36199_80 | 0.90 | 73.26 | 0.0000 | 0.0028 | Winter minimum phosphate concentration |
| 50000_74 | 0.65 | 14.83 | 0.0049 | 0.0089 | Winter minimum phosphate concentration |
| 58479_12 | 0.62 | 13.25 | 0.0066 | 0.0089 | Winter minimum phosphate concentration |
| 18407_279 | 0.68 | 17.25 | 0.0032 | 0.0089 | Annual phosphate concentration range |
| 18407_279 | 0.64 | 14.14 | 0.0055 | 0.0089 | Spring phosphate concentration range |
| 50000_74 | 0.59 | 11.65 | 0.0092 | 0.0095 | Spring phosphate concentration range |
| 36199_80 | 0.67 | 16.59 | 0.0036 | 0.0089 | Winter phosphate concentration range |
| 50000_74 | 0.61 | 12.75 | 0.0073 | 0.0089 | Winter phosphate concentration range |
| 18407_279 | 0.66 | 15.45 | 0.0043 | 0.0089 | Winter phosphate concentration range |
| 63214_51 | 0.70 | 18.53 | 0.0026 | 0.0089 | Annual mean silicate concentration |
| 63214_51 | 0.86 | 48.04 | 0.0001 | 0.0056 | Winter mean silicate concentration |
| 61897_186 | 0.76 | 24.95 | 0.0011 | 0.0074 | Winter mean silicate concentration |
| 1701_190 | 0.68 | 16.86 | 0.0034 | 0.0089 | Winter mean silicate concentration |
| 16142_260 | 0.61 | 12.73 | 0.0073 | 0.0089 | Winter mean silicate concentration |
| 63214_51 | 0.70 | 18.76 | 0.0025 | 0.0089 | Annual maximum silicate concentration |
| 63214_51 | 0.77 | 26.77 | 0.0008 | 0.0074 | Winter maximum silicate concentration |
| 50460_263 | 0.62 | 13.16 | 0.0067 | 0.0089 | Winter maximum silicate concentration |
| 16449_9 | 0.74 | 22.97 | 0.0014 | 0.0074 | Annual minimum silicate concentration |
| 63214_51 | 0.71 | 19.62 | 0.0022 | 0.0089 | Annual silicate concentration range |
| 63214_51 | 0.77 | 26.33 | 0.0009 | 0.0074 | Winter silicate concentration range |
| 50460_263 | 0.65 | 14.61 | 0.0051 | 0.0089 | Winter silicate concentration range |

p-value _adjusted_ are based on a Benjamini-Hochberg correction (Benjamini & Hochberg 1995)

Table S 3 Characterization of high-quality BLASTx.

Matches obtained in comparison of *E*. *maclovinus* SNP against NCBI database. We only retained SNPs located in genes with putative functions that are compatible with the hypothesis of local adaptation.

| SNP | Detection method | Gene | Species | Protein name | e-value | Hit length | General function |
| --- | --- | --- | --- | --- | --- | --- | --- |
| 44791_149 | fsthet | slc39a6 (210632) | *Parambassis ranga* | zinc transporter ZIP6 isoform X2 | 1,712E-29 | 739 | MF: metal ion transmembrane transporter activity |
| 6291_214 | LFMM | slc8a2a (8218) | *Gymnodraco acuticeps* | sodium/calcium exchanger 2a | 2,000E-20 | 914 | MF: calcium: sodium antiporter activity; calmodulin binding; metal ion binding  BP: cell communication |
| 42_59 | LFMM | vtg3 (8218) | *Gymnodraco acuticeps* | Phosvitin | 6,131E-21 | 1266 | MF: lipid transporter activity; nutrient reservoir activity |

MF: Molecular function

BP: Biological process
